# Engineering a microfluidic-assisted 3D bioprinting approach for the hierarchical control deposition and compartmentalisation of graded bioinks

**DOI:** 10.1101/2025.02.10.637484

**Authors:** Federico Serpe, Lucia Iafrate, Marco Bastioli, Martina Marcotulli, Caterina Sanchini, Valeria De Turris, Michele D’Orazio, Biagio Palmisano, Arianna Mencattini, Eugenio Martinelli, Mara Riminucci, Carlo Massimo Casciola, Giancarlo Ruocco, Chiara Scognamiglio, Gianluca Cidonio

**Affiliations:** Department of Mechanical and Aerospace Engineering, University of Rome “La Sapienza”, 00184 Rome, Italy; Center for Life Nano- & Neuro- Science – CLN^2^S, Italian Institute of Technology (IIT), 00161 Rome, Italy; Department of Electronic Engineering University of Rome Tor Vergata, Rome, Italy; Department of Molecular Medicine, University of Rome “La Sapienza”, Rome, Italy

**Author notes:** Corresponding Authors: Dr Gianluca Cidonio, Dr Chiara Scognamiglio.

**Keywords:** bioprinting, microfluidics, gradient, cell patterns, interface

## Abstract

The advent of 3D bioprinting has revolutionised tissue engineering and regenerative medicine (TERM). Today, tissues of single cell type can be printed with extreme resolution and printing fidelity. However, the ultimate functionality of the desired tissue is limited, due to the absence of a multicellular population and diversity in micro-environment distribution. Currently, 3D bioprinting technologies are facing challenges in delivering multiple cells and biomaterials in a controlled fashion. The use of interchangeable syringe-based systems has often favoured the delamination between interfaces, greatly limiting the fabrication of interconnected tissue constructs. Microfluidic-assisted 3D bioprinting platforms have been found capable of rescuing the fabrication of tissue interfaces, but often fails to guarantee printing fidelity, cell density control and compatimentalisation. Herein, we present the convergence of microfluidic and 3D bioprinting platforms into a new deposition system capable of harnessing a microfluidic printhead for the continuous rapid fabrication of interconnected functional tissues. The use of flow-focusing and passive mixer printhead modules allowed for the rapid and dynamic modulation of fibre diameter and material composition, respectively. Cells were compartmentalised into discrete three-dimensional layers with defined density patterns, confirming the punctual control of the presented microfluidic platform in arranging cells and materials in 3D. *In ovo* and *in vivo* studies demonstrated the functionality of 3D bioprinted constructs with patterned vascular endothelial growth factor (VEGF) and transforming growth factor-β1 (TGF-β1), respectively. This, in turn, facilitated the simulation of diverse cellular environments and proliferation pathways within a single construct, which is currently unachievable with conventional 3D bioprinting techniques, offering new opportunities for the fabrication of functionally graded materials and physiologically-relevant skeletal tissue substitutes.

## Introduction

Over the past decade, biofabrication strategies have advanced beyond current technological frontiers to surpass the limitation in tissue three-dimensional assembly [1,2]. A crucial challenge in 3D bioprinting functional tissue substitutes is to simultaneously control cellular density, biomaterial properties and resulting microarchitecture, to engineer functionally-graded 3D constructs [3,4]. Indeed, the spatial control over the composition of extruded inks on-demand increases the heterogeneity of 3D bioprinted constructs, which is a crucial aspect for the engineering of a physiological microenvironments. Particularly, when dealing with tissue interfaces, replicating the gradual variation of biological, mechanical, and chemical properties is fundamental to mimic the native environment with elevated accuracy [5]. This is currently challenging to obtain with conventional 3D bioprinting approaches. To reproduce tissue interfaces, extrusion-based 3D bioprinting platforms typically rely on the use of multiple extruders to switch between bioinks with different material compositions [6] or cell types [7]. Besides the complex set-up required to house multiple syringes, the resulting 3D structures are fabricated in separate compartments for each tissue portion, lacking a continuous and gradual variation, with consequent poor multi-tissue functionality [8–10].

The coupling of a microfluidic-based extruder with 3D bioprinting systems (*microfluidic-assisted 3D bioprinting*) has come to the fore as a breakthrough technology in recent years to offer a possible solution to the engineering of functionally-graded materials and tissues [11–15]. Currently, microfluidic printheads consist of a modular system where a microfluidic chip is connected to a coaxial nozzle (typically fabricated from glass capillaries or coaxial needles) that functions as the extruder [16,17]. The ionic crosslinking instantly occurs at the tip of the nozzle, where the ionic solution meets the gel precursor. This convenient approach enables the decoupling of the spinning process from the biomaterial rheology, allowing the extrusion of low-viscosity biomaterial inks [18]. However, the presence of physical constraints (*i.e.* needles or capillaries) limits the control over fibre characteristics and raises the amount of shear forces localised at the extrusion tip, which is detrimental for encapsulated cells and bioink deposition [19,20]. Moreover, manual assembly of the coaxial nozzle often results in frequent operational problems, influencing the coaxial centring of one aperture within the other, which may prevent the high-throughput printing of 3D scaffolds due to recurrent nozzle clogging. Lastly, the resulting printing fidelity and functionality are often unable to drive the fabrication of functionally-graded tissue constructs [15]. Despite extensive progresses in building a connector-free monolithic microfluidic printhead, many challenges concerning the extrusion precision and the size of the 3D bioprinted tissues remain open. A new powerful approach (chaotic 3D assembly) has been recently introduced as an alternative to coaxial-wet spinning and 3D bioprinting [21] to offer the unparalleled possibility to pattern micro-architectural features within a single extruded fibre. However, the technology is not yet scalable and often reduced to the fabrication of fibres, with limited yield for the assembly of hierarchical scaffold architectures. Flow-focusing-based microfluidic printheads have been proposed to assemble wet-spun fibres in three-dimensional constructs [22–24]. However, evident limitations in terms of layer stacking and three-dimensional assembly have been reported, demonstrating the unsolved technical control of the flow-focusing approach. Commercially-available microfluidic-assisted 3D bioprinting platforms have enhanced the biofabrication process by stacking hybrid, hollow or core-shell fibres controlled by pneumatic valve technology that allowed for the modulation of the fibre size and material switching [25]. Nevertheless, the need for a porous vacuum substrate to continuously drain the crosslinking solution outflow limited the printing resolution and fabrication possibilities [26,27]. During the last decade, a number of microfluidic printheads have been employed to fabricate both simple and core-shell fibres for applications in neural [28,29], renal [30] and muscular [31] modelling. However, despite the increased functionality of printed constructs, both the high survival rate even for high-density cell printing and the generation of well-organized 3D constructs were not always accomplished, limiting the range of possible applications.

In this work, we present a novel strategy to harness newly designed microfluidic printing heads able to spin microfibres with tuneable characteristics, enabling the assembly of hierarchically-organized 3D tissue models via the unprecedented spatial control of biomaterials, cells and biomolecules. The need to insert external components such as needles or glass capillaries is eliminated, further streamlining the fabrication process and minimizing possible cellular damage during fibre extrusion. Following microfluidic chip designing and printing validation, the possibility to realize three-dimensional structures with functional patterning of biomaterial inks and cells is here investigated. For the first time, we reported the 3D bioprinting of functionally-graded constructs where cell density can be adjusted on-demand with high spatial control. Leveraging the tuneable logic of the newly designed microfluidic printheads, this study confirms the possibility to engineer hierarchical scaffolds where cells are compartmentalized in separate yet interconnected microenvironments, creating biological niches within the same 3D structure. This is further demonstrated for the engineering of functionally-graded bone constructs by spatially patterning human bone marrow stromal cells (HBMSCs) to elicit a functional differentiation towards skeletal lineage. Pre-clinical investigation of functionally-graded skeletal constructs demonstrated an improved capacity of cellular implants incorporating key biological factors to drive vascular infiltration and skeletal tissue formation *in vivo*. This novel approach enables the production of functional scaffolds in one step that could potentially drive encapsulated cells towards different proliferation pathways, moving a step forward in replicating native tissue interfaces.

## Materials and Methods

### Microfluidic chip fabrication

A printhead master mold containing the negative of the desired geometry was used to shape polydimethylsiloxane (PDMS) into the final printhead. CAD design was processed via a 3D editor (NAUTA^®^+) and 3D slicing software for printing on an XFAB 3500SD SLA printer (DWS, Italy). Printed molds were washed with absolute ethanol for 10 min followed by UV and thermal curing for 24 h. PDMS (SYLGARD™ 184, Dow Corning, USA) was prepared by mixing the base component with curing agent in a ratio 10:1, cast on the mold and stored in oven for 2 h at 75 °C. Following PDMS removal, access ports to internal microchannels were created with 1.5 mm puncher. Two negatively-cast PDMS slabs are joined together via air plasma treatment for 2 min.

### Microfluidic chip design optimization

A microfluidic chip was engineered to serve as the 3D bioprinter extrusion system. The flow-focusing (FF) module was optimized to allow continuous spinning of fibres and guarantee correct fibre extrusion and deposition by minimizing sheath overflow. The FF microfluidic printhead is equipped with a monolithic PDMS nozzle and microchannels with a diameter of 100 μm. The micromixer (MM) printhead integrated a new passive micromixer upstream of the flow-focusing junction, which consisted of an adapted version of a previously developed Tesla mixer [32].

### Biomaterial composition and preparation

A nanocomposite material ink was prepared from a blend of 0.5% w/v Laponite^®^ (XLG grade, BYK Additives & Instruments, UK), 2% w/v alginate (Mw ≈ 33 kDa, Protanal 1740, FMC Biopolymers) and 1.5% w/v gelatin (type A, bloom 300, from porcine skin, Sigma-Aldrich, UK) as previously reported [33]. Briefly, all powders were sterilized under UV light for at least 30 min, prior to inclusion in stirring deionized water (DW) for overnight suspension and solubilisation. In the case of cell embedding, the final biomaterial ink – referred to as LAG – was exposed to UV light for additional 30 min and kept at 40°C. A crosslinking solution was prepared from 330 mM CaCl_2_ in 80% glycerol (Sigma-Aldrich, UK) following through dispersion. The final solution was lastly sterile-filtered and kept at 4°C until usage.

### Bioprinting set-up

A custom-made 3D bioprinter machine was coupled with a syringe pump system composed of a base and three low-pressure moduli (neMESYS, CETONI GmbH, Germany). Syringes were loaded with inks and the crosslinking solution and plugged to the microfluidic printhead through plastic Tygon^®^ tubes (Saint Gobain, France). Printed scaffolds were fabricated with a typical fibre distance of 1mm, dimensions ranging from 6×6 to 20×20 mm and layering of 10, 20 or 30 layers with an axial step of 100 μm. After the printing procedure, samples are dipped in sterile 330 mM CaCl_2_ solution for 5 min to complete crosslinking. FF device was used for printing single-material scaffolds, while the MM printhead was adapted for the printing of multi-material constructs.

### Characterization of the microfluidic printheads

#### Fibre diameter

To measure the fibre diameter on a chip, flowrates were varied between 25, 30, 35 and 40 μl/min for the core and from 20 to 50 μl/min with 5 μl/min steps for the sheath flow. To visualize the biomaterial ink flow under an inverted fluorescence microscope (AX10, Zeiss, Germany), fluorescein-tagged alginate (FITC-alginate) was used in the blend. After a settling time of 30 seconds to allow flow stabilization, images were acquired with an integrated camera (Axiocam 305 color, Zeiss, Germany) and processed with ImageJ software. Similarly, fibre diameter has been characterized after printing. Fibres were deposited onto a 1 mm-thick microscope glass slide at a constant speed of 12 mm/s in a linear pattern and flowrates were changed according to the aforementioned conditions. Then, core and sheath flowrates were kept at 30 μl/min each and the feed rate of the printer was varied from 9 to 15 mm/s. Images of deposited fibres were acquired and processed with ImageJ software.

#### Micromixer

The MM chip has been characterised in terms of the duration and quality of the mixing. Two LAG compositions were adapted to contain either FITC-alginate or untagged alginate. The MM device was placed on the microscope and real-time images were acquired at the output of the micromixer. Following the non-FITC solution flow stabilization at 30 μl/min, the fluorescent solution was flown at 30, 20, 10 and 5 μl/min. After the switching, images were acquired every 5 sec for the first 30 sec, then every 10 sec until 2 min. Images acquired every 2 minutes were used to evaluate the quality of the mixing by calculating the *mixing index* (MI) parameter [34,35], whose formula is here reported:

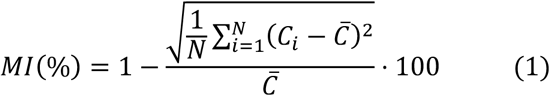

where *N* is the number of investigated points in the region under analysis, *C_i_* is the pixel intensity of the point under investigation and 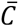 is the mean pixel intensity of the entire investigated region. Finally, the MI was calculated when keeping the same flowrates for FITC and non-FITC solutions but progressively increasing their intensity, reaching an overall sum of 10, 15, 20, 25, and 30 μl/min.

### Mass loss and swelling properties

Mass loss and swelling ratio were measured according to previous work [36]. Briefly, mass loss analysis was carried out on samples obtained either by material casting (bulk, n=3) or 3D-printed in a grid shape (3DP, n=3). The initial weight of each sample (*m*_0_) was calculated after crosslinking the samples and carefully drying the latter with a paper sheet. Samples were then incubated in Hank’s balanced salt solution (HBSS, Sigma-Aldrich, UK) at 37°C and removed every 3-4 days. After being carefully dried with a paper sheet, samples were weighted (*m_i_*) and submerged in fresh HBSS.

The percentage of weight loss was calculated as:

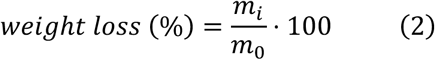

Swelling analysis was performed on bulk LAG to investigate the difference in water uptake when 165 mM and 330 mM CaCl_2_ solutions were used for crosslinking. LAG bulk samples were weighted after crosslinking (*m*_0,*sol*_) to calculate the initial wet mass and then lyophilized to obtain the dry mass (*m*_0,*dry*_). These two components allow to calculate the macromer fraction as:

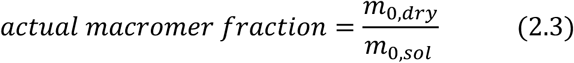

The other samples were incubated in HBSS at 37°C and collected at time points (*i* = 2, 4, 6, 24 h) and weighted before (*m*_0,*sol*_) and after (*m*_0,*dry*_) lyophilization. The sol fraction was calculated as:

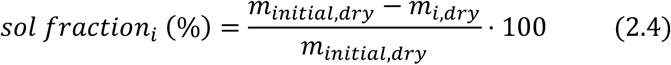

Where *m_initial,dry_* = *m*_0,*sol*_ · *actual monomer fraction*. The mass swelling ratio *q* at time point *i* was instead obtained as:

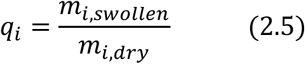

where *m_i_*_,*swollen*_ = *m_i_*_,*sol*_ – *m_i_*_,*dry*_.

### Cell culture

#### MG63 cell culture

MG63 osteosarcoma cells (ATCC, USA) were cultivated in DMEM high glucose cell culture medium (Sigma-Aldrich, USA) supplemented with 10% (v/v) fetal bovine serum (FBS), 1% (v/v) Penicillin/Streptomycin (Merck Life Sciences) and 1% L-glutamine (Merck Life Sciences), and kept in culture at 37°C with 5% of CO_2_, changing media every 3-4 days.

#### Human bone marrow stromal cell culture

Human Bone Marrow Stromal Cells (HBMSCs) from healthy donors were used following the Declaration of Helsinki and its later amendments and approved on June 22, 2023 by the Institutional Review Board (Department of Molecular Medicine, Sapienza University of Rome, Italy). HBMSCs were isolated as previously described [37]. Briefly, bone fragments were collected as surgical waste from healthy subjects undergoing orthopaedic surgery. Bones were scraped and minced into small pieces in order to release bone marrow cells. After vigorous pipetting, cell suspensions were collected in Minimum Essential Medium with alpha modification (Merck Life Sciences, Saint Louis, USA) supplemented with 20% (v/v) FBS (Thermo Fisher Scientific, Waltham, USA), 1% (v/v) Pen/Strep and 1% L-glutamine. Cells were seeded at the density of 2 × 10^5^ cells/cm^2^ and grown at 37°C and 5% CO_2_ balanced air. The next day, the culture medium was replaced to remove non-adherent cells and HBMSCs were grown until confluence. Cells were expanded for a maximum of three passages and then transplanted following 3D bioprinting.

### 3D printing of hierarchical acellular scaffolds

To visualize the layered deposition of multiple materials, 28-layer structures were printed, sectioned in the axial direction and tilted at 90° to visualize the biomaterial patterning in the axial plane. FITC-alginate and untagged material ink were modulated with variable flow rates to print 3D structures with alternate, step and gradient constructs. A confocal laser-scanning microscope (Olympus IX83, USA) was employed to acquire images that were subsequently analysed with ImageJ. To demonstrate the ability to tailor the width of the interface between different materials, FITC-alginate LAG and rhodamine-tagged LAG were loaded in different syringes and connected to the input of the MM chip. Assuming as 100% the total printing time of the scaffold, different switching set-ups were performed preserving the overall flowrate of 30 μl/min. More specifically, considering the quantity solution1/ratio1:1/solution2, we printed scaffolds with 50/0/50%, 42.5/15/42.5% and 30/30/30% compositions.

### 3D bioprinting of hierarchical cell-laden scaffolds

Cells were collected from culture flasks and pelleted prior to labelling with lipophilic tracers (Vybrant^®^ DiD, *646-663 nm* or DiO, 4*83-501 nm* cell-labelling solution, ThermoFisher Scientific, USA) following the manufacturer’s protocol and as previously reported [38]. Briefly, cells were resuspended at a concentration of 1×10^6^ cells/mL in a serum-free medium adding 5 μl of Vybrant^®^ dye for each mL and incubated in the dark (37°C and 5% CO_2_) for 20 minutes. Afterwards, the cell solution is centrifuged, and the supernatant is removed and replaced with clean medium at least 3 times to ensure the removal of any unbounded particle dye. For the generation of a single-cell gradient, the MM chip was used to combine LAG with and without MG63 cells. The cells-laden LAG contained Vybrant^®^ DiD-labelled MG63 at a concentration of 15×10^6^ cells/ml. Flowrate ratios between the two syringes were varied from 0 to 1 at intermediate steps of 1:6, 1:3, 1:2, 2:3 and 5:6, maintaining the overall sum at a fixed value of 30 μl/min. The resulting gradient allowed for precise variation in cell density across the scaffold. The number of cells was estimated for each condition through ImageJ software, considering as 100% the number of cells present in the sole cell-laden material. For scaffolds with increasing cell-type concentrations, MG63 cells were labelled with two distinct fluorescent dyes (Vybrant® DiD and DiO) and resuspended at a concentration of 10 × 10⁶ cells/mL. Cell suspensions were mixed within the MM chip, where the ratio of flow rates was modified between consecutive print runs. This approach yielded two-layer scaffolds with uniform but distinct cellular architectures. For all conditions, each cell type is normalized against the overall number of cells counted for each flowrate condition. For the fabrication of thick hierarchical scaffolds, 24-layer structures were printed maintaining an overall flowrate of 30 μl/min. MG63 cells were marked with Vybrant^®^ DiD and resuspended in LAG at a concentration of 15 x 10^6^ cells/mL and loaded in a syringe. Cell-free LAG material was loaded in a separate syringe. Alternate, step and gradient 3D scaffolds were realized and imaged as aforementioned. Acquired images were analysed with ImageJ software. For the generation of multi-cellular interface scaffolds, 24-layer structures were printed, and the flowrate switching was performed as reported in previous sections. The scaffolds were then imaged from (i) the bottom, (ii) the top, and (iii) the side, after cutting longitudinally and tilting the 3D construct at 90°. Acquired confocal images were processed with ImageJ software and assembled in mosaics.

### Viability of cell-laden scaffolds

Cell viability was investigated on HBMSCs on day 1 and day 7 using Calcein AM (C1430, Invitrogen, Thermo Fisher Scientific) staining and acquisition via confocal microscopy. At the fixed time point, cells were washed three times with HBSS and incubated at 37°C and 5% CO_2_ in serum-free media with Calcein AM at a concentration of 0.6 μg/ml in a dark incubator for 1 hour. Afterwards, samples were washed 3 times with HBSS and then imaged with a confocal laser scanning microscope (Olympus IX83, USA). Cell viability was quantified using ImageJ software as previously reported [38,39]. Briefly, cells stained both with Calcein and DiD were considered alive, while cells stained only with DiD were considered metabolically inactive (dead). The percentage of viability is derived from the ratio between Calcein-positive cells and DiD-stained cells.

### Alkaline phosphatase (ALP) staining on 3D cell-laden scaffolds

ALP staining was carried out following previously employed protocols [33,40]. Briefly, control (no cells) or HBMS-laden scaffolds with 1 or 3 x 10^6^ cell/mL were washed thrice with HBSS, fixed with 95% ethanol solution for 10 mins, allowed to dry in air for 1h and washed twice with HBSS. ALP solution was prepared from Naphtol AS-MX Phosphate and Fast Violet Salt (Sigma-Aldrich, UK) in DW and used to stain the 3D constructs during incubation for 1h at 37°C. Samples were then rinsed with DW and imaged with an inverted microscope (AX10, Zeiss, Germany) in transmission mode.

### Immunostaining of cell-laden scaffolds

After fixation with 4% paraformaldehyde (PFA)/HBSS for 30 min at RT, 3D printed samples were washed with HBSS and then permeabilized with 0.5% Triton X-100/HBSS for 1h at RT. After 3 washes in HBSS, samples were blocked with 3% BSA (BSA, Sigma-Aldrich) in HBSS for 1h at RT. For actin staining, 1h autofluorescence blocking in 1% (w/v) BSA is carried out prior to the addition of AlexaFluor488 phalloidin dye (Excitation/Emission (nm) 495/518, Invitrogen, UK), followed by HBSS washing steps and imaging. For immunofluorescence staining, primary antibodies (rabbit Osteocalcin, Bioss antibodies, 1:200; mouse Osteopontin, Invitrogen, 1:300; rabbit Collagen 1, Bioss antibodies, 1:200) were incubated in 1% BSA in HBSS overnight at 4°C. Samples were then washed with HBSS 0.1% Tween 20, and incubated with Alexa Fluor™ fluorophore-conjugated secondary [40,41]antibodies (Invitrogen, 1:400) in 1% BSA in HBSS, Alexa Fluor™ 488 Phalloidin (Invitrogen) and Hoechst (Sigma-Aldrich, for nuclei visualization) for 1h at RT. Samples were then washed with HBSS 0.1% Tween 20 and stored at 4°C in HBSS.

### Chick chorioallantoic membrane (CAM) assay (*in ovo*)

Samples were 3D printed using LAG ink with (VEGF-loaded) or without (control) vascular endothelial growth factor (VEGF, ImmunoTools, Germany). The highest concentration of VEGF was selected based on compound localisation demonstrated in previous publications [42]. By modulating VEGF between 0 and 100 μg/ml with a gradient approach (VEGF-gradient), hierarchical patterning was achieved. Animal procedures were strictly regulated by European Directive 2010/63/EU and by Italian Legislative Decree No. 26 dated 4th March 2014. At Day 0 chicken eggs were incubated in a Hatching incubator (CT120SH, Cimuka, Turkey) for 14 days at 37.7 °C in a 50% humified atmosphere while turning 90° every 3.2 h. On day 7 of incubation, a scalpel was used under sterile conditions to create a 1 cm^2^ window on the eggshell in order to place the treatment groups over the CAM. A sham control (empty) was implemented in this study. Eggs were sealed with sterile parafilm secured with tape, incubated without rotation and daily checked. On Day 14 of incubation, scaffolds were harvested and CAM integration was inspected and imaged with a Zeiss Stemi 305 stereomicroscope with a digital camera. Subsequently, the gestational process was terminated under specific guidelines.

### AI-assisted analysis of vasculature

To quantify the afferent vessels with an automated approach, we employed previously developed AI-assisted analysis [33]. Briefly, the dataset was assembled from images of CAM vessels acquired with a Zeiss Stemi 305 stereomicroscope at 1x magnification. Three images for each of the four treatment groups were acquired. The analysis was carried out in four steps as follows: vessel segmentation, area and diameter computing, and Chalkley score (CS) calculus.

#### Vessel segmentation

Matlab 2024^®^ was used to carry out image analysis. Segment Anything Model (SAM) [43] was employed to perform the vessel segmentation, with foreground and background points as prompts. The final outputs of the segmentation step consist of binary images with true (false) values in pixels belonging (not belonging) to vessels. *Bwlabel* function was then used to separate (in the 8- connection sense) and identify the single vessels. To mitigate potential bias in the analysis of the empty class, a central region corresponding to the average area of a bioprinted construct was removed, effectively setting it to the background.

#### Area and diameter computing

The area was identified by recurrent sum of the area of the non-zero pixels identifying the specific object, i.e., vessel. To compute the diameter of the single vessels the *bwdist* function (which computes for each pixel the distance from the nearest non-zero pixel) was applied. To determine the centre of the vessel, we skeletonized the image using the *bwskel* function, which ensured that the central value corresponded to the radius. To exclude vessel endings, values below the 5th quantile of the distribution were discarded.

#### Chalkley’s score computing

Chalkley’s score (CS) was determined within an interactive framework. To centre and scale the Chalkley grid, the user is asked to draw a bounding box containing the implanted construct. The number of intersections between circles of the Chalkley grid and at least one vessel is counted at different rotation angles from 0° to 360° with step 10°. Only the ten highest CS values were considered for each image.

### Subcutaneous implantation and histological analysis

Cell-free control and cell-laden constructs were dosed with loaded TGF-β1 and transplanted subcutaneously into the back of a 2-month-old female CB17.Cg-Prkdscid Lystbg-j/Crl (SCID/beige) mice (Charles River, Wilmington, USA). After 4 weeks, samples were harvested, fixed in 4% formaldehyde for 48h and decalcified in 0.5M EDTA for 7 days. Samples were then processed for paraffin embedding following standard procedure. Four-micron-thick sections were stained with haematoxylin and eosin (H&E) for morphology evaluation, and with Sirius red to highlight collagen deposition.

### Statistical analysis

GraphPad Prism 8 (GraphPad Software Inc. La Jolla, CA) was employed for statistical analysis. Data variation was assessed by the D’Agostino-Pearson normality test. Obtained results were evaluated by one-way ANOVA with a Tukey’s multiple comparison test with a single pooled variance and considering significative difference if p < 0.05.

## Results and Discussion

The microfluidic-assisted 3D bioprinting technology here proposed wants to advance beyond current limitations in extrusion-based bioprinting approaches, aiming to control in space cells, material inks and biomolecules. By harnessing a new microfluidic printhead, a hierarchical biological compartimentalisation can be achieved following 3D deposition. Cells and biomolecules are included within the fibre prior to deposition and precisely patterned in 3D with a single-step process, avoiding delamination and facilitating different cell-type communication and multi-tissue maturation (**Figure 1**).

**Figure 1.**
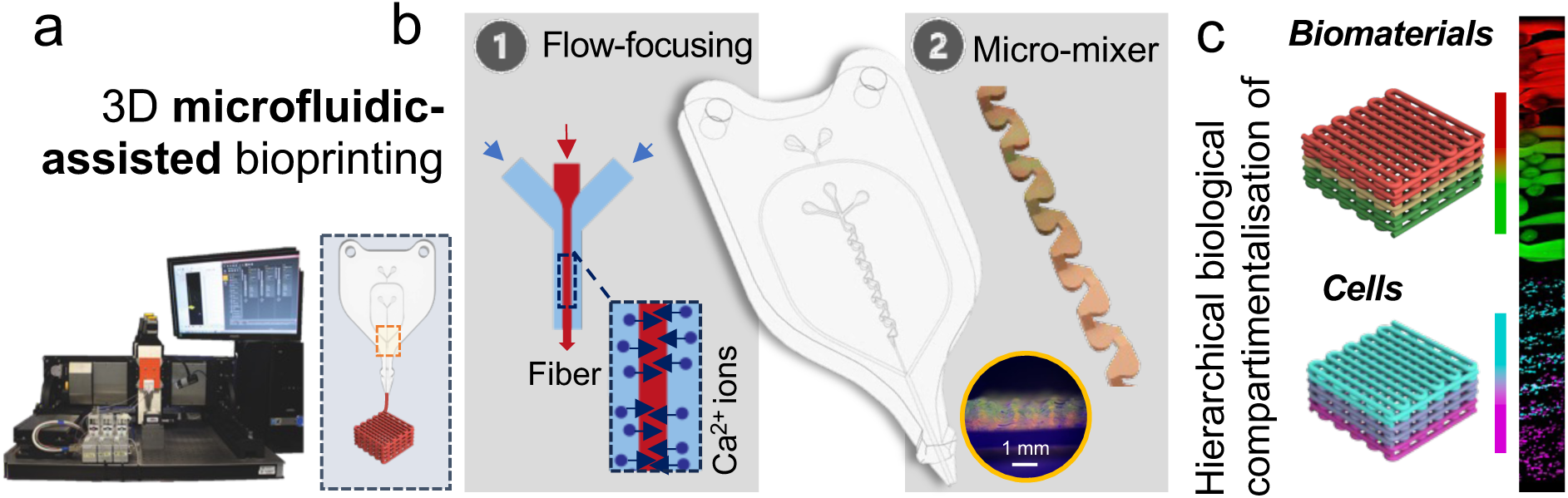
Microfluidic-assisted 3D bioprinting for the fabrication of hierarchically organized 3D constructs. a) The 3D bioprinting system is equipped with a 3-axis motorized machine to drive the movement of the microfluidic printhead and a syringe pump system to control the input flowrates independently. b) The fibre production is facilitated by the sheath flow of Ca^2+^ ions in the crosslinking solution, which diffuses in the biomaterial ink promoting crosslinking via the flow-focusing junction. The integration of a micromixer structure creates turbulence within microchannels allowing to homogenise the input components. According to the relative flowrates of the two bioinks, the output blend will contain a variable percentage of the two input fluids. c) The microfluidic-assisted 3D bioprinting system allows spatiotemporal control over deposition, enabling the realization of hierarchically organized constructs of simple biomaterials or bioinks (material ink + cells). Heterogeneous 3D structures can be realized with a single-step process and high control, dramatically increasing the complexity and biomimicry of printed scaffolds.

### Microfluidic-assisted spinning allows the tuning of fibre diameter and deposition resolution

The engineered microfluidic printheads were obtained via PDMS-casting of SLA printed moulds and assembled to avoid the use of metallic or glass nozzles, considerably decreasing manufacturing complexity and variability. This approach resulted convenient in terms of cost and time with respect to conventional photolithography, representing a valuable alternative to photolithography for microfluidic fabrication [44,45]. The flow-focusing microfluidic printhead (FF chip, **Figure 2a**) was fabricated and integrated with a custom-built 3D bioprinting system [46] enhancing the resolution of the printed 3D constructs (**Movie S1**, **Figure S1**). A previously developed nanocomposite material (Laponite^®^ - alginate - gelatin, LAG [33]) was employed to investigate printability and skeletal tissue functionality. This composite ink has been found ideal for coaxial-extrusion [33], thus resulting an ideal candidate for deposition with the FF printhead. Considering the bioactive nature of nanoclay and gelatin materials, alginate is here employed as synergistic functional support for both mineralisation and rapid crosslinking [40,41]. Depending on the relative flow intensity of the input components, the composition of the extruded fibre is adjusted in real-time and on-demand. FF microfluidic printhead was designed and implemented for the printing of single-core fibres. By adjusting fluid flowrates with tuneable microfluidic pumps, the range of printed fibre diameter was investigated (**Figure 2b,c**). The resulting diameter of fibres produced in a flow-focusing-based channel configuration depends on the relative intensity of the core and sheath flowrate imposed [47], according to the formula reported in **Figure 2b**. By selecting unique flowrates for the core flowrate, the sheath flowrate was changed to accommodate the values that would allow for subsequent deposition of the extruded fibre on a substrate. We called this printability range.

**Figure 2.**
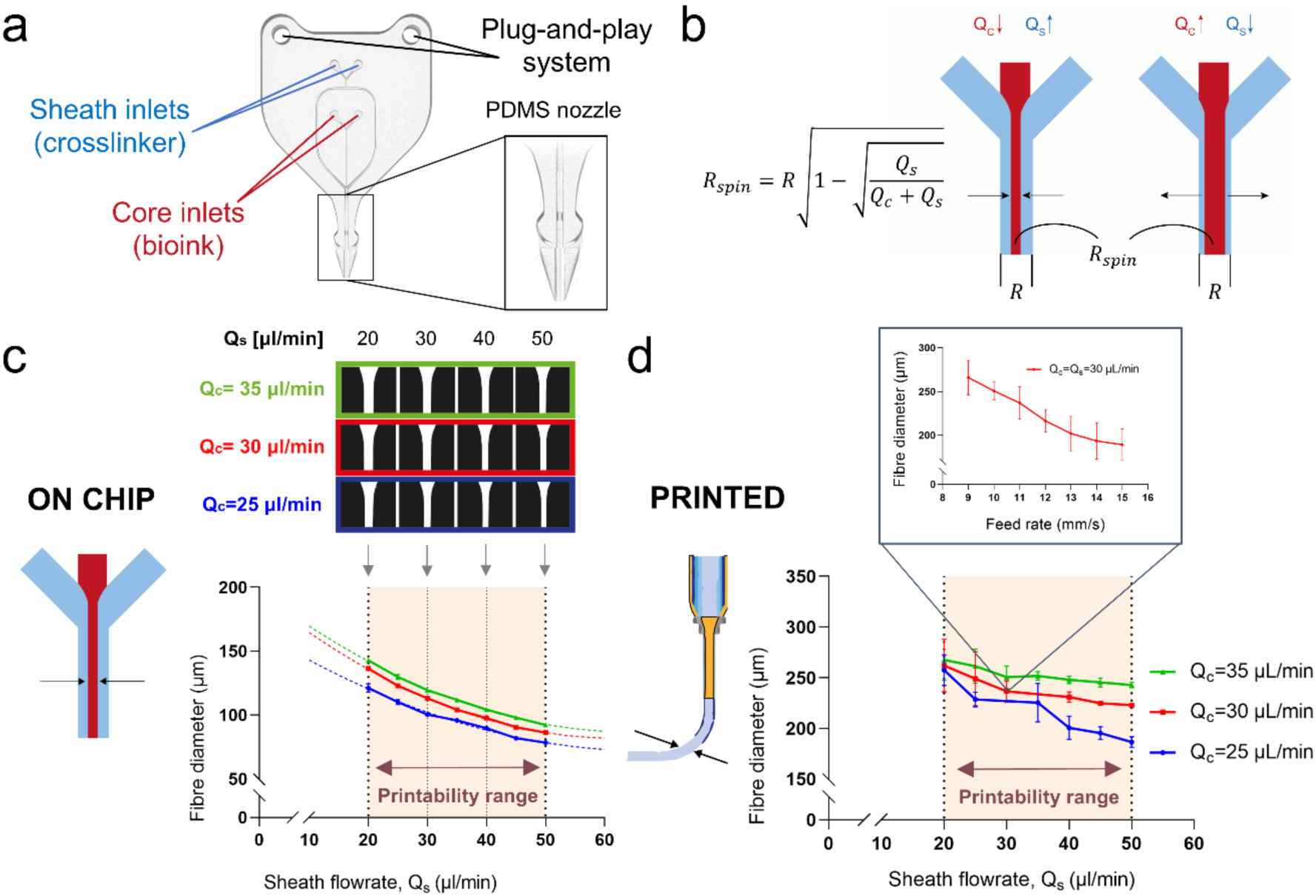
FF chip design and properties. a) 3D rendering of the FF chip with an enlarged view of the PDMS nozzle. b) Sketch illustrating fibre dimensions when varying the core and sheath flowrates, showing a thinner fibre when either the core flowrate is decreased, or the sheath flowrate is increased and a larger fibre for the opposite conditions. The formula on the left describes the behaviour of the fibre radius, where *R* is the radius of the channel and *R_spin_* the radius of the spinned fibre. c) Characterization of the fibre diameter on-chip. In the upper part, processed images of the fluorescent core material reveal the variation in the width of the central flow for different flow conditions. d) Characterization of the deposited fibre diameter in function of different flowrate conditions. In the inset, the fibre diameter was calculated for fixed flowrates and variable feed rate to show the dependence from printing speed. Results are expressed as mean ± SD of at least three replicates for each experiment.

On-chip fibre diameter was characterized and an exponential decay behaviour was observed, due to increased shearing force applied by the sheath fluid (**Figure 2c**). For each core flowrate, the resultant fibre diameter can be adjusted of ± 50 μm, allowing to control the core flow diameter continuously from a minimum of 78 ± 1 μm to a maximum of 155 ± 2 μm. The measured diameters result smaller compared to the one reported by previous attempts where the fibre sizes ranged between 200 and 800 μm [22]. In a further study, Beyer and co-workers reported the possibility to control fibre diameter from 75 to 300 μm [48]. However, in the latter case, the possibility to deposit fibres with a such large difference in diameter was enabled by the presence of a porous printing substrate connected to a vacuum source to drain the excess sheath fluid that hampered good resolution of the final structure. The effective diameters of deposited fibres at a fixed printing velocity of 12 mm/s were measured (**Figure 2d**). The results confirmed a linear decrease of the fibre diameter with increasing the sheath flowrates, consistent with the on-chip measurements. The observed overall increase in the deposited fibre diameter compared to the on-chip filament was likely attributed to the material crosslinking mechanism. Following the extrusion process, deposited fibres could be observed to vary in diameter from 186 ± 5 μm to 337 ± 3 μm. By fixing core and sheath flowrates to 30 μl/min (in an optimal deposition condition) the diameter of the deposited fibres was characterized for different feed rates (**Figure 2d**). Deposited fibres were observed to vary of 70 μm in diameter (from 189 ± 18 μm to 266 ± 20 μm) when the printing velocity was diminished from 15 to 9 mm/s, respectively. Previously reported attempts with metal-based coaxial nozzles demonstrated that, by varying the deposition speed from 6 to 1 mm/s with a fixed flowrate of 5 μl/min, the fibre diameter was increased from 150 μm to about 300 μm [17]. The greater range of attainable spun fibre diameters is possibly due to the presence of a coaxial needle as the outlet, which required lower printing speed and flowrates, resulting in diameters increasingly susceptible to variations of the pulling force.

### 3D printing acellular scaffolds with high precision and reproducibility

Following the investigation of fibre formation within the printhead chip, the assessment of 3D construct fabrication was carried out. FF printhead allowed the deposition of scalable three-dimensional lattice structures. This was demonstrated by printing increasingly larger structures with 6, 10 and 15 mm^2^ area, respectively (**Figure 3a**). By tuning core and sheath flowrates, 3D scaffolds were found to be printable at high-resolution and high-throughput consistency (**Figure S2a**). Deposited fibres could be layered in 3D with a fine control allowing for the spatially-resolved printability both at 90° (**Figure S2b-i**) or 45° (**Figure S2b-ii**) stacking.

**Figure 3.**
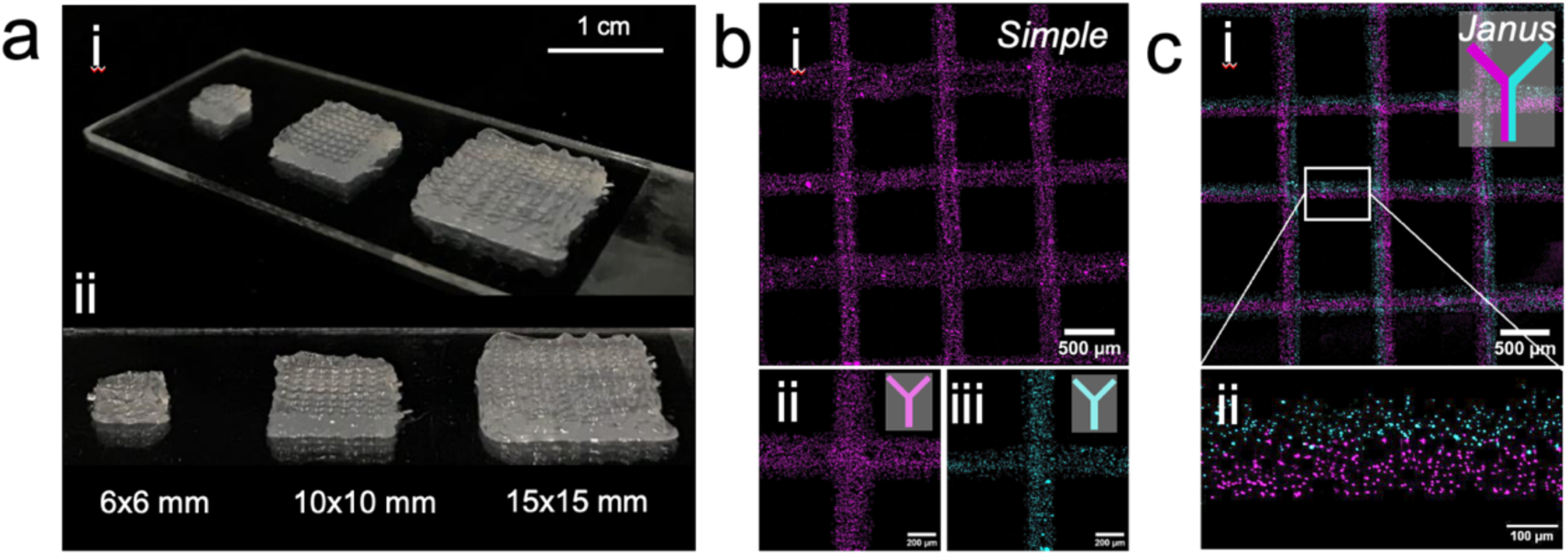
Microfluidic printing of simple 3D scaffolds. a) Printing of acellular grid scaffolds with scalable dimensions, ranging from 6×6 to 20×20 mm. Confocal view of 3D deposition of microfluidic-spun fibers. b) Printing of grid scaffolds with single materials loaded with fluorescent beads. A larger view (i) and single crossing (ii and iii) of the 3D printed scaffolds. c) Printing of grid scaffolds with two materials loaded with fluorescent beads of different colours. A larger view (i) and close-up on a single fiber to show the hybrid composition (ii).

The FF chip printhead allowed the control over the extrusion of simple (**Figure 3b**) or Janus (**Figure 3c**) fibres whether one or two materials were input. Particularly, this approach allowed the hosting of different cell types in close proximity. Such a strategy has been employed in numerous reported works in the TERM field [49–51], especially for cardiovascular [17] and muscular [52] regeneration. To investigate the stability of deposited fibres and whether the printing process affected the rate of material degradation, acellular 3D-printed scaffolds were maintained in culture (37°C, 5% CO_2_) for up to 21 days and compared to casted (bulk) controls (**Figure S3a**). Bulk and 3D printed scaffolds were found to lose a non-significantly different amount of weight over time up to 14 days. However, following culture for 7 more days bulk scaffolds were observed to hold 37% of the original weight, while the printed constructs continued to degrade until reaching 5% of the initial weight. This behaviour was probably due to the greater area/volume ratio of grid 3D construct with respect to the bulk control condition that implied faster degradation rates [41]. Despite the high degradation of 3D printed constructs measured, these results are not detrimental, as it is hypothesised that cells embedded within the fibres would naturally deposit ECM over time, contributing to the reinforcements of the 3D printed construct over time. Finally, in order to determine the influence of the crosslinker concentration on the chosen biomaterial ink formulations, swelling measurements were carried out. Specifically, the percentage of soluble fraction and the mass swelling ratio (q) were assessed in samples with different crosslinking percentages (**Figure S3b,c**). When a less concentrated calcium buffer (165 mM CaCl_2_) was used, uphold water uptake facilitated the release of uncrosslinked ink and the swelling of the crosslinked portion of the material. In contrast, samples crosslinked with 330 mM CaCl_2_ solution exhibited 10% less swelling but showed a greater soluble fraction due to the reduced swelling capacity. This swelling behaviour may prevent the scaffold from preserving the macrostructure, as excessive water absorption may alter the resulting mechanical properties [53,54]. Based on these findings, the crosslinker concentration of 330 mM was identified as optimal for the crosslinking of cellular 3D constructs.

### Microfluidic mixer enables to control the ink composition in real-time

Inspired by a simplified version of the Tesla mixing design [32], the micro-mixer (MM chip) printhead was equipped with a convoluted three-dimensional geometry upstream to the flow-focusing junction (**Figure 4a-i**). Two input materials were blended in a variable ratio before reaching the flow-focusing junction, where they were solidified in a fibre shape and extruded (**Figure 4a-ii,iii**). The ultimate fibre composition was adjusted on-demand by controlling the flowrate intensities. To characterize the mixing process, two key factors were considered: (i) the *mixing duration*, defined as the time required by the system to produce a stationary flow following a change in flow rates; (ii) the *mixing quality*, considered as the degree of homogenization of the two input solutions at the mixer outlet. Dynamic observation of the mixer output (**Figure 4b**) revealed that for any of the analysed combinations, 120 s is a sufficient time to consider flow stabilization after switching to a different flow condition. For every investigated flowrate, a minimum of 60 s was found to be required to arrive at 90% of the complete mixing. The mixing index (MI) provides an absolute measure of mixing performances and was calculated following an elapsed time (120 s) after switching flow rates from 0 μl/min to 5, 10, 15, 20, 25 and 30 μl/min (**Figure 4c**). The results show that mixing quality increases with larger flowrates differentials while decreases for low-intensity switching. This is because lowering the flowrate intensity generates less turbulence within the microchannels, leading to a reduction in the efficiency of the mixing process. In addition, the MI of two solutions forced at the same flowrate has been evaluated here (**Figure 4d**) finding a gradual and significant (p<0.05) increment in MI when the overall sum of the flowrates is increased. This phenomenon is again explained by the increase in turbulence (quantified via the Reynolds number) that rose linearly with the fluid velocity. For high flowrate values, thus, stochastic mixing is encouraged, resulting in increased performances of the mixer. The values obtained for the MI are in line with the ones found in the literature for biofabrication mixing applications, supporting the efficacy of this approach for precise and efficient mixing [11]. It is worth to highlight that the passive mixing unit here presented for the first time is based on a simplified geometry adapted from a Tesla mixer [32]. The MM chip is equipped with mixing units lacking a flow-splitting feature for ease in fabrication and improved flow stability before the flow-focusing junction. Active mixing unit would ameliorate the fast mixing between ink solutions, improving the mixing index [55]. However, a passive micro-mixer design was here adapted due to the ease in fabrication and the possibility to control the compact dimensions in the limited confined space of the microfluidic printhead.

**Figure 4.**
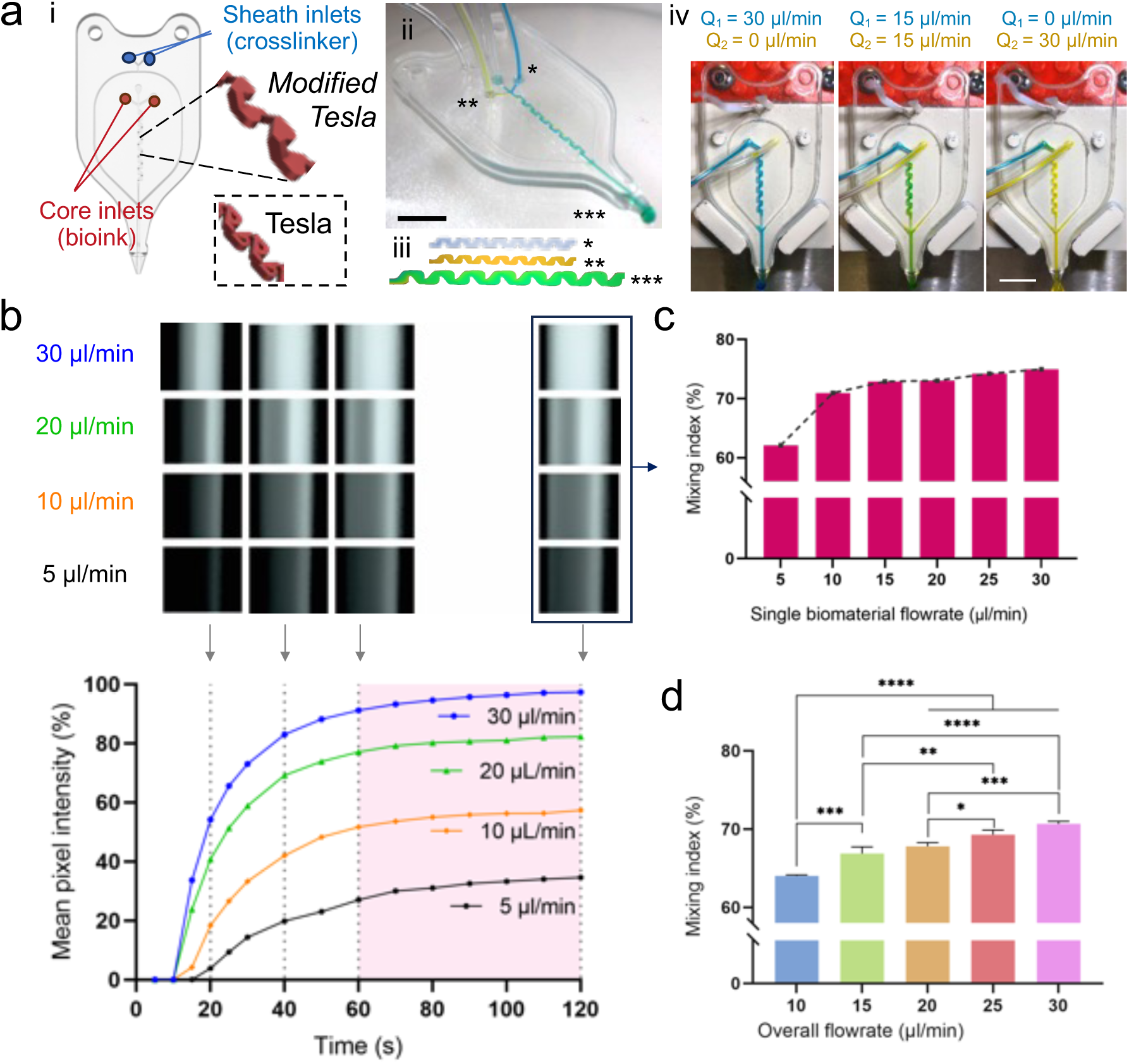
Comprehensive analysis of the MM chip. a) 3D rendering i) of the MM chip with detailed modified Tesla geometry compared to the canonical Tesla units. ii) Macrograph of the MM printhead iii) with a close-up of the micromixer geometry with unaltered and mixed inks. iv) In-situ macrographs of the MM chip during printing when yellow- and blue-coloured solution is inserted in input and green output are observed. Depending on the flowrate ratio, the output colour is different. Scale bars are 1 cm. b) Characterization of the microfluidic mixer via dynamic analysis of the mixing process. In the graph, the mean pixel intensity during time is reported to visualize the dynamic behaviour of the material after switching. In all these cases, a settling time of 60 s was required to reach 90% of the final value. In the upper part, images were acquired at the output of the microfluidic mixer at different time points (20, 40, 60 and 120 s) for various flow conditions. c) Calculation of the mixing index 120 s after switching. On the x-axis, the flowrate of the FITC material after switching is reported. d) Calculation of the mixing index when varying flowrate intensities. In this case, the x-axis represents the sum of the two flowrates (FITC and untagged material) showing a significant augmentation for increased flowrates. Statistical significance was calculated via one-way ANOVA. Results are expressed as mean ± SD of at least three replicates for each experiment, *p<0.05, **p<0.01, ***p<0.001, ****p<0.0001. Scale bars: a-ii,iv, 15 mm.

However, MI results significantly lower with respect to other passive micromixers geometries [56–58] and in particular to Tesla-based designs, in which MI is reported to reach up to 94% [32] with considerably shorter mixing length. This was due to the simplified Tesla structure here proposed, in which the fluid splitting and recombination are eliminated for the practical arrangement of the printhead channels. Herein, mixing tests were carried out with hydrogel-precursor materials (LAG), which exhibit higher rheological properties with respect to water as previously reported by our group [33]. This factor required an increase in the physical length of the micromixer in order to ensure adequate mixing performances.

### Patterning of biomaterial inks in 3D constructs

Given the opportunity to modulate the composition of extruded materials on-demand, we investigated the ability of the proposed MM printing head to control the patterning of biomaterial inks in 3D. More specifically, biomaterial density profiles (**Figure 5a**) as well as multi-material interfaces (**Figure 5b**) can be readily obtained with tailored arrangements. The final architecture of the obtained 3D constructs recalled the one typically observed in human tissue interfaces (e.g., osteochondral interface), which often include mechanical, physical and chemical modification in the extracellular environment [5]. Alternate (**Figure 5a-i**), step **(Figure 5a-ii)**, and gradual (**Figure 5a-iii**) variations of the FITC-tagged ink with respect to the clear ink on the transversal plane were observed and quantified. From acquired confocal images, defined regions with specific biomaterial ink arrangements were successfully obtained, confirming the possibility to reproduce pre-defined 3D patterns along the axial direction. To validate the possibility to assemble compartments with spatially-controllable interfaces, the material ink tagged either with FITC or rhodamine was independently dosed, switching from one to another during printing (**Figure 5b**). The control over the spatial patterning of the interface width was confirmed by microscope observation, resulting in the effective realization of 3D interfaces with on-demand width, which can range from 250 up to 1000 μm.

**Figure 5.**
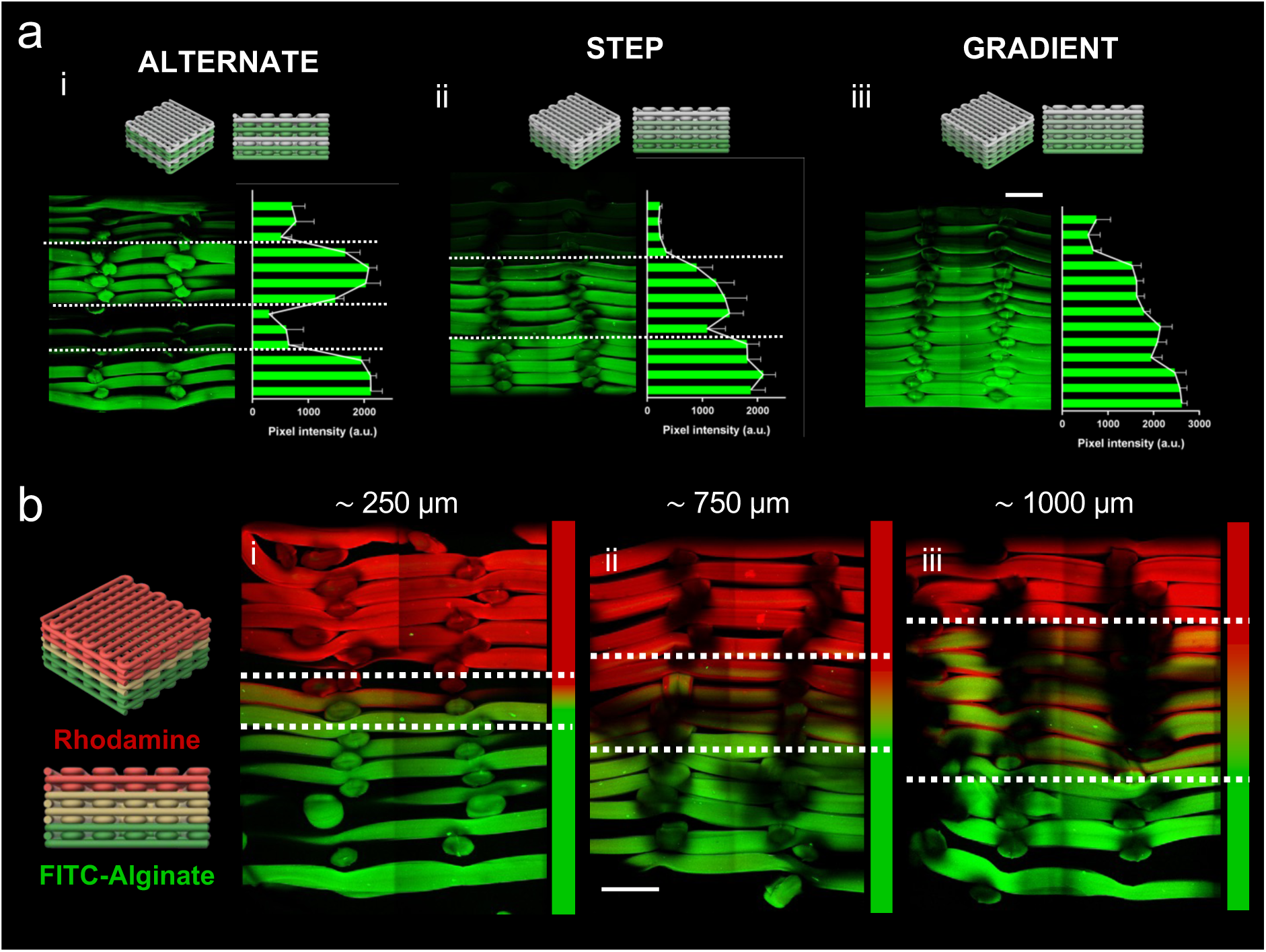
3D patterning of materials. a) Layered arrangement of LAG material containing FITC-tagged and normal alginate to visualize the variation of the signal intensity in the transverse plane. Alternate, step and gradient variation of the composition was obtained by adjusting the flowrates of the FITC and NON-FITC material. Images were taken at the confocal microscope after cutting the scaffolds along the axial direction and tilting each half of 90°. Results are expressed as mean ± SD of at least three regions chosen inside the fibres. The scale bar is 200 μm. b) Realization of tailored material interfaces within the 3D scaffold using LAG enriched with fluorescent dyes (FITC and rhodamine). Confocal images of the transverse section of the scaffolds revealed the formation of an interface (i.e. the area where the fibre contained a mixture of the two materials) that can be varied from 250 to 1000 μm. Scale bars are 200 μm.

### Compartimentalisation of cells in 3D scaffolds

To investigate whether the proposed microfluidic system could adjust cell density in real-time, MG63 cells were printed with varying densities (**Figure 6a-i**). By modulating the ratio between the acellular and the cell-laden ink flowrates via the microfluidic printhead, cell density could be controlled during printing (**Figure 6a-ii**). The post-printing quantification of the cell density percentage (**Figure 6a-iii**) showed great fidelity with respect to the expected value, which was given by the ratio between the two flowrates intensities. To further demonstrate the enhanced control over cellular deposition, the modulation of cellular composition during printing using two cell groups is here reported. Cells were marked with two cell-labelling dyes (**Figure 6b-i**) and printed to create scaffolds with controlled density transitions. The density ratio between the two flowrates was gradually switched from 0 to 1 (**Figure 6b-ii**) so that a smooth variation was observed in the overall cell population of the scaffolds. Following deposition, the quantification of the cell density based on tagging fluorophores revealed the accurate deposition and spatial arrangement obtained with the relative flowrate ratio (**Figure 6b- iii**).

**Figure 6.**
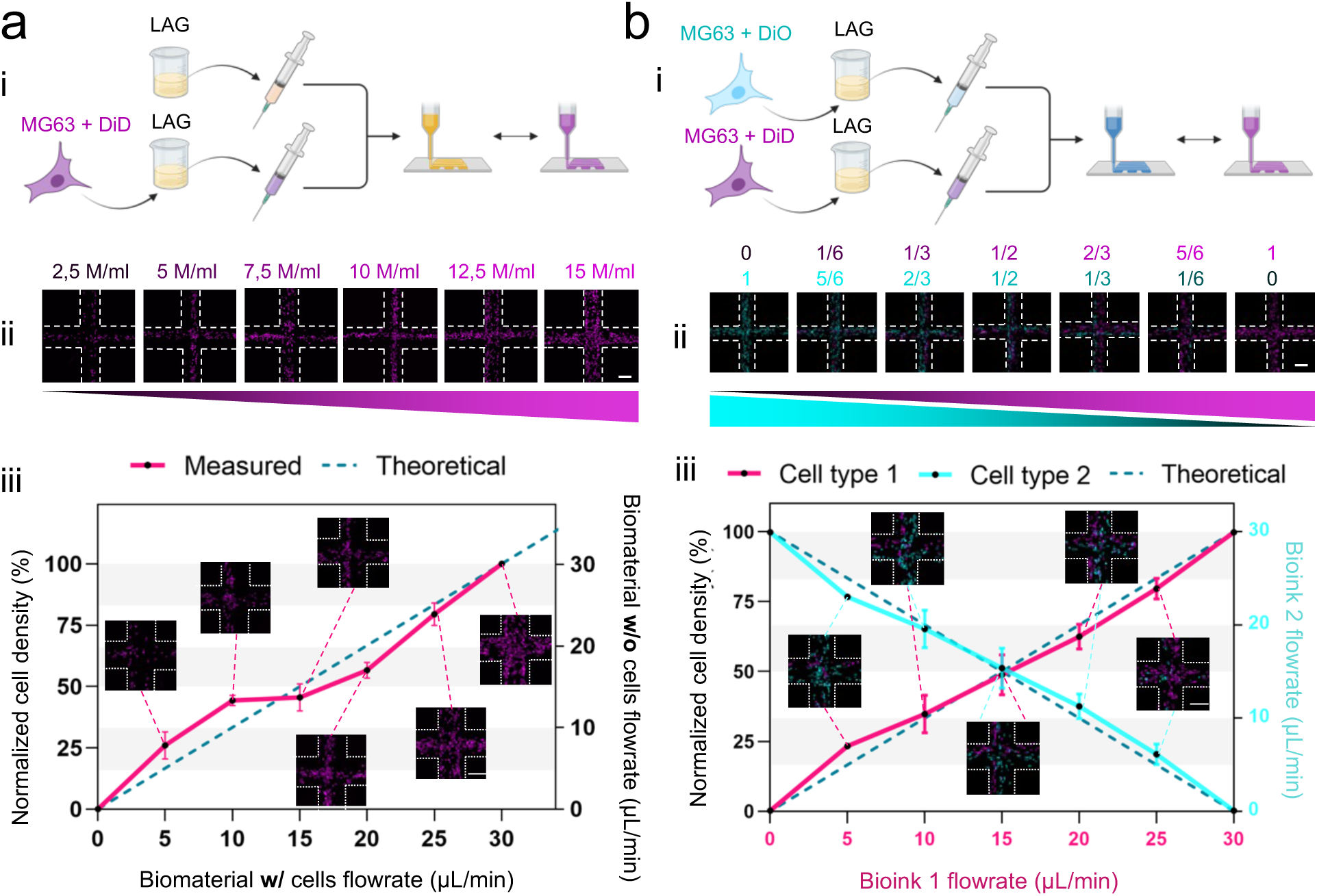
Cell gradients. a) Cell density variation of fluorescently labelled cells. i) A cell-laden and a cell-free ink are prepared and separately input in the micromixer. ii) Confocal images of 3D printed scaffolds with variable cell numbers and (iii) quantification of the number of cells for each flowrate condition. b) Cell density variation of MG63 cells marked with different fluorescent dyes. i) The two bioinks are separately connected to the micromixer. ii) Confocal images of the 3D printed scaffolds with variable types of cell populations and (iii) quantification of the number of cells belonging to the two pre-selected groups. In both cases, the dashed line represents the expected number of cells of each condition. Results are expressed as mean ± SD of at least three replicates for each experiment. Scale bars are 200 μm.

This methodology demonstrates significant potential for hierarchically patterning cells in layered interfacial constructs. To validate the approach, a cell-laden LAG ink was dynamically modulated in flowrate during volumetric deposition. This precise control enabled the fabrication of distinct cell density profiles, including alternate (**Figure 7a-i**), stepwise (**Figure 7a-ii**), and gradual (**Figure 7a- iii**) variations. A microfluidic-based mixing approach was employed by Idaszek and collaborators to recreate the interface between hyaline and calcified cartilage with a 3D-printed model [11]. Similarly, Kuzuku and co-workers proposed a 3D bioprinting system equipped with a static mixer to control cellular density in 2D and in 3D [59]. However, the ability to modulate the interfacial region width and density was not reported before. To investigate the possibility of generating spatially-controlled interfaces, living cells labelled with differently-emitting dyes were simultaneously extruded and modulated using the custom micromixer (**Figure 7b**). This approach allowed the serial deposition of either cyan (Channel 1) or magenta (Channel 2) cells with pre-defined densities. Confocal images demonstrated no trace of cross-talking between the differentially compartmentalised cell populations. The axial patterning was confirmed by the orthogonal view (**Figure 7c**) clearly highlighting the graded cell distribution. By altering the switching conditions, interfaces of increasing dimensions (up to 1000 μm) were engineered. These results confirmed the ability of the microfluidic printhead to tailor the cell distribution in 3D with exceptional spatial control, a critical feature for accurately replicating tissue interfaces and advancing the design of biomimetic constructs.

**Figure 7.**
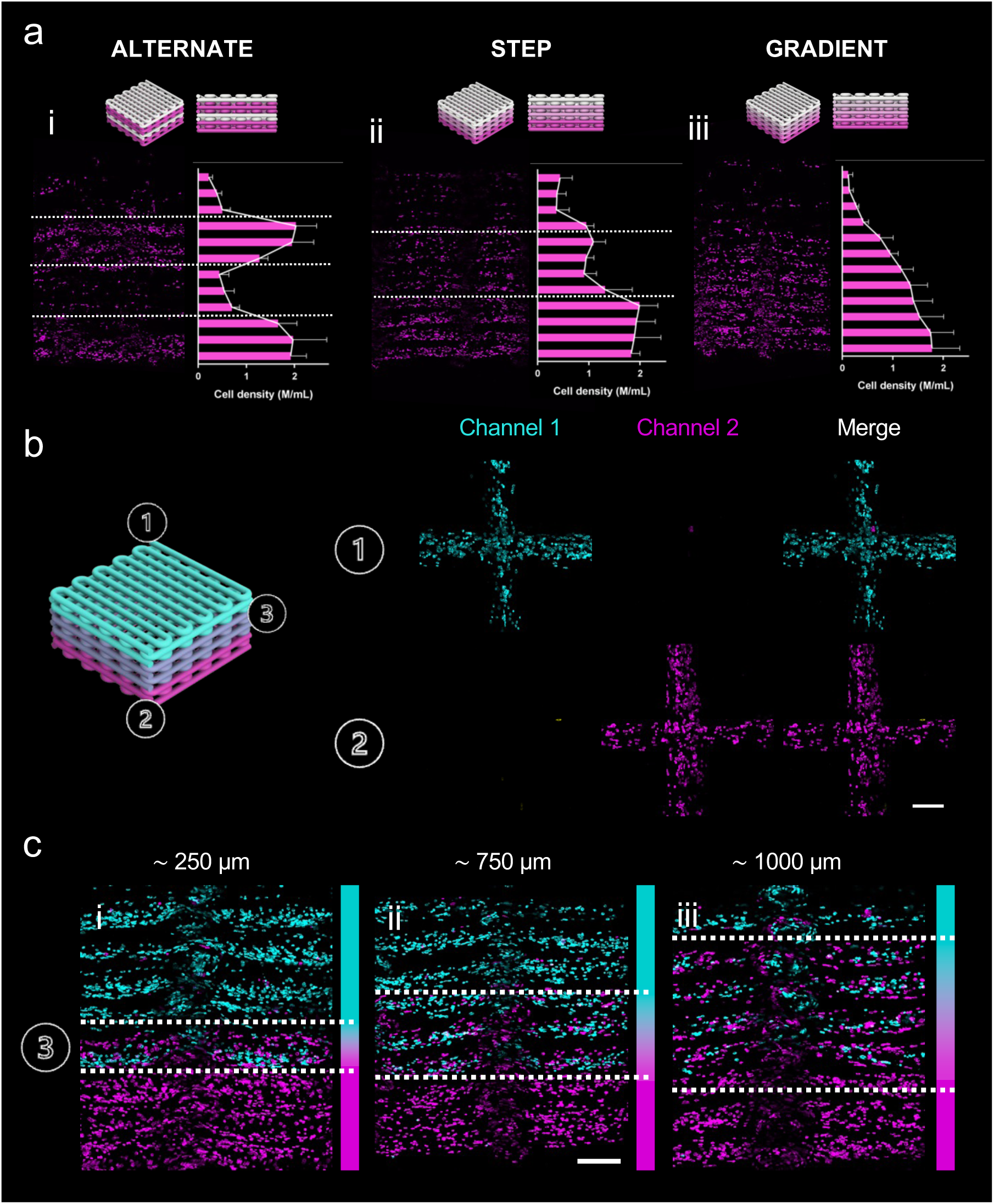
3D patterning of cells. a) Hierarchical arrangement of MG63 cells labelled with fluorescent tracer within 3D printed scaffolds. Alternate, step and gradient concentration of cells was observed and quantified after cutting the scaffold along the axial direction. b) 3D patterning of two cell groups (MG63 marked with different fluorescent tracers) to realize 3D scaffolds with variable interface widths. Images were taken (1) from the top, (2) from the bottom and (3) from the side. In the latter case, three different widths of the interface portion, i.e. where two cell families coexist, are reported. The interface area can be adjusted between 250 and 1000 μm with high control. Results are expressed as mean ± SD of at least three replicates for each experiment. Scale bars are 200 μm.

### Microfluidic 3D bioprinting can pattern skeletal stem cells while preserving viability and supporting functionality

To investigate the possibility to control cellular density prior to differentiation, we encapsulated and patterned in 3D HBMSCs using the microfluidic-based printhead approach. The viability of 3D bioprinted constructs was analysed following 1 (**Figure 8a-i, ii**) and 7 (**Figure 8a-iii, iv**) days of culture, with no significant difference with casted (bulk) constructs (**Figure 8b-i**), confirming the mild conditions for shear and bioprinting delivery with the microfluidic-based approach. These results confirm that the microfluidic-assisted 3D bioprinting process preserves cell viability and proliferation. Concomitantly, 3D bioprinted cells were found to demonstrate a sustained proliferation in spatially-organized construct. We hypothesize that the diffusion of nutrients was more favoured in grid-like 3D structures which hold a higher surface area to volume ratio (*i.e.* a larger area of the biomaterial is exposed to the medium). Moreover, the overall high cellular viability obtained post-printing results consistent with the theoretical evaluation formulated in previous publications, highlighting the protective effect of the sheath flow over the core spun fibre [14]. In accordance with this assumption, ongoing studies will be carried out to further elucidate the protective mechanism provided by the microfluidic printhead, potentially offering a significant advantage in maintaining cell integrity during 3D bioprinting.

**Figure 8.**
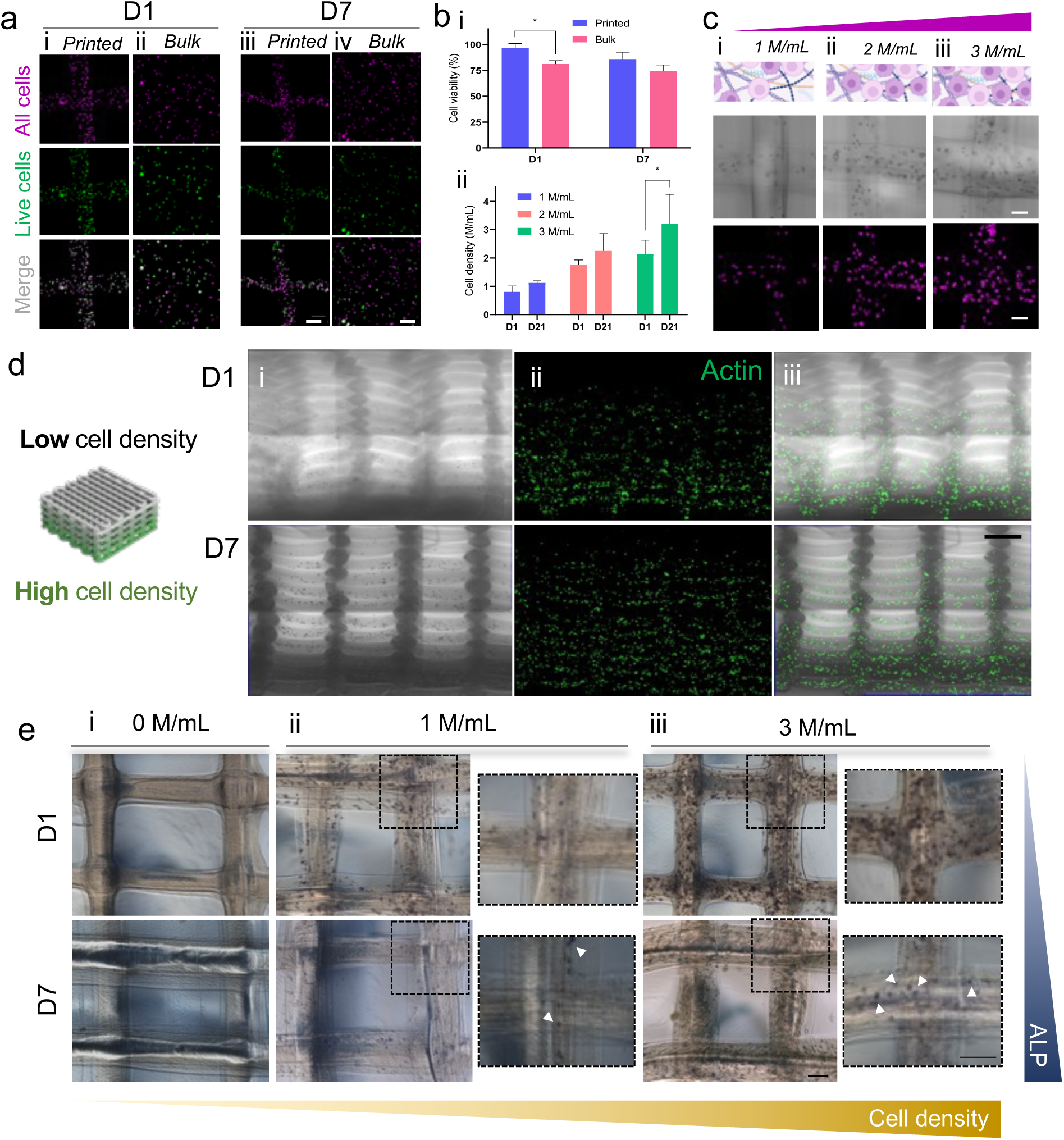
Bioprinting of HBMSCs. a) Viability analysis of HBMSCs comparing printed scaffolds to bulk bioink. Cell viability was quantified as the ratio between Calcein-stained cells to the total number of cells, which are stained with lipophilic tracer (i). The scale bar is 200 μm. In the graph b) the quantification of cell viability. c) Bioprinting of 3D constructs with different densities of HBMSCs (i), ranging from 1 to 3 million cells per ml. d) A single construct fabricated with varying cell concentration along the orthogonal direction with low (1 M/mL) up to high (3 M/mL) cell densities, is investigated for cell distribution from 1 up to 7 days. e) Macrographs and close-up of ALP staining of (i) cell-free, (ii) 1 M/mL and (iii) 3 M/mL cellular densities. The scale bar is 200 μm. Statistical significance was calculated via one-way ANOVA. Results are expressed as mean ± SD of at least three replicates for each experiment, *p<0.05.

The microfluidic-assisted 3D bioprinting approach here presented was adapted to selectively deposit 1, 2 or 3 million cells per volume of ink (M/mL) to monitor the proliferative capacity of HBMSCs up to 21 days. Results demonstrated the significant (p<0.05) proliferative ability of bioprinted HBMSCs exclusively at 3 M/mL (**Figure 8b-ii**). However, a lack of correlation between the intended and the actual printed cell densities was observed, particularly at higher concentrations. This discrepancy is likely due to the progressive sedimentation of cells within the bioink precursor during the printing process. The use of a mixing system within the syringe dispenser could eventually solve this discrepancy. The ability to print specific cell densities was confirmed microscopically (**Figure 8c**) with both brightfield and confocal micrographs, demonstrating a lower cell-to-cell distance associated with increasing printing density. Notably, the enhanced density of printed cells fostered greater cell-to-cell interactions potentially contributing to a controlled enhancement of skeletal fate commitment in the overall functional properties of the construct. This highlights the potential of precise cell density modulation to influence cellular behaviour and optimize the functional outcomes of bioprinted tissues. HBMSCs were printed in a graded process from high (3 M/mL) to null (0 M/mL) cellular density (**Figure 8d**). Following 7 days of culture, resulting constructs were found stable with a confirmed proliferative capability and ultimate increase in cell number, visible following actin staining (**Figure 8d-i, ii, iii**). Thus, HBMSCs were 3D bioprinted with increasing densities to investigate the skeletal differentiation following osteogenic media culturing over 7 days (**Figure 8e**). Results illustrated that following the controlled deposition of a greater density (3 M/mL), ALP was found highly expressed following 7 days of culture compared to a lower (1 M/mL) concentration. The increase in ALP expression correlated with a greater cell density possibly due to increased cellular communication as previously demonstrated [60]. The use of this new microfluidic-assisted 3D bioprinting technology allowed for the compartmentalisation of cell densities to investigate the functional effect of graded patterning over skeletal differentiation of bioprinted HBMSCs. Thus, the functional investigation of HBMSCs 3D bioprinted with a graded approach focused on the higher-density regions of the interfacial constructs. This analysis revealed enhanced clusterisation of Collagen I (**Figure S4a**) as well as increased levels of Osteocalcin and Osteopontin (**Figure S4b**) expression at day 7 of culture in osteogenic media, confirming the findings related to the optimal functional response of HBMSCs printed at 3 M/mL density.

### In ovo and in vivo implantation confirmed the functionality of graded 3D constructs

To demonstrate the spatial control over the segregation of biomolecules in 3D, both *in ovo* and *in vivo* assays were carried out. Functionally-graded 3D bioprinted constructs were implanted in a CAM model to investigate vascular-induced growth via graded VEGF patterning. Empty (**Figure 9a-i**) and factor-less (**Figure 9a-ii**) controls were prepared and compared with VEGF-gradient (**Figure 9a-iii**) and adsorbed (loaded, **Figure 9a-iv**). Quantitative evaluation of the number of afferent vessels (Chalkley Score, **Figure 9b-i**) revealed significantly (p<0.01) augmented vascularisation of functionally-graded 3D constructs compared to VEGF-free control, with no evident improvement with respect to the adsorption of VEGF. No changes were identified in the vascularised area among all groups (**Figure 9b-ii**). This was possibly due to the low VEGF concentration used in the study, to imitate CAM response to physiological doses. Nevertheless, the diameter of afferent vessels was found significantly (p<0.01) enhanced in VEGF groups compared to empty control, with a greater effect identified in VEGF-loaded (p<0.05) compared to VEGF-graded constructs. The rapid response of VEGF biologic is well-known in literature and here anticipated to enhance local angiogenesis due to the nanoclay-mediated adsorption of VEGF compared to the graded inclusion via microfluidic-assisted 3D bioprinting [40].

**Figure 9.**
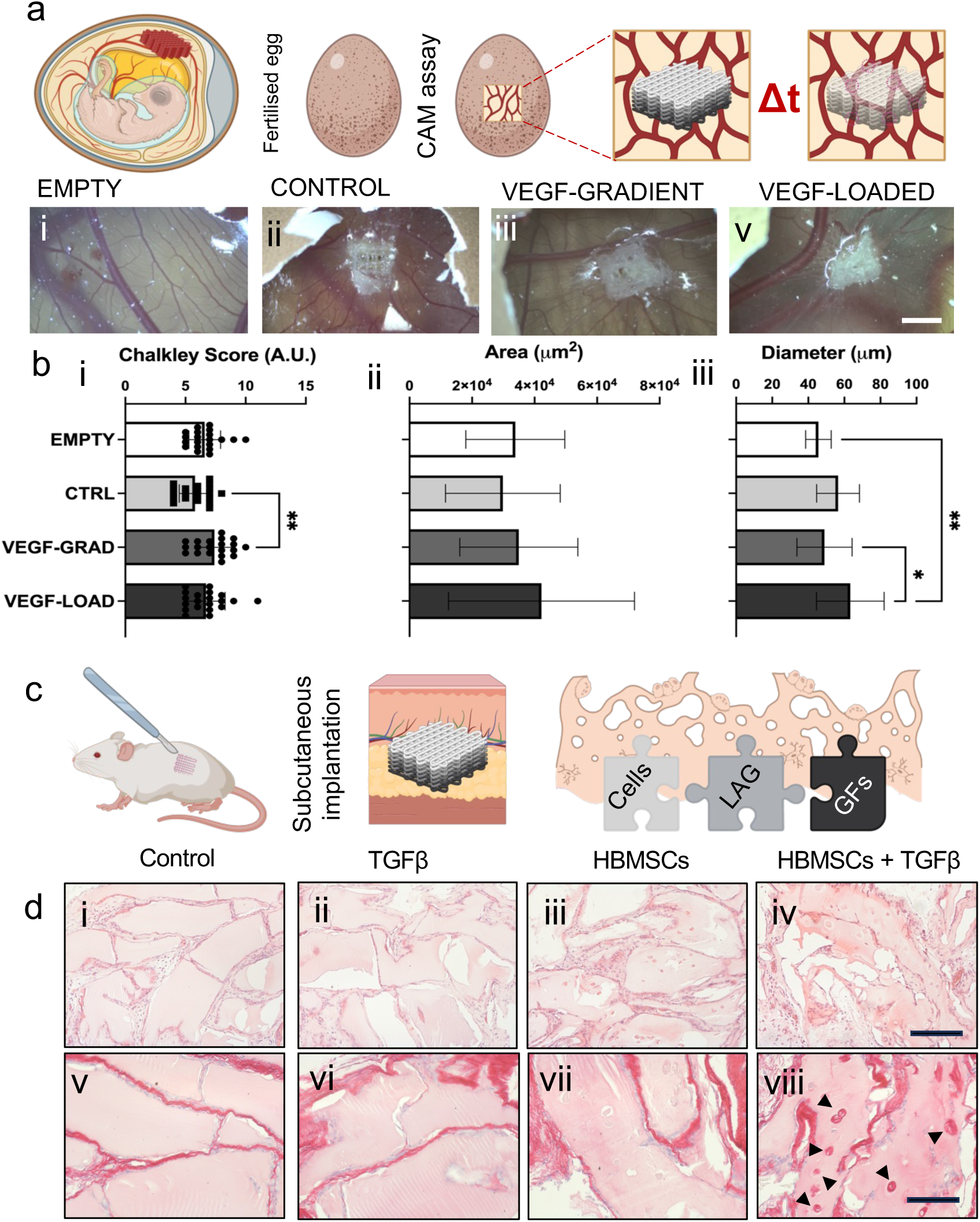
In ovo and in vivo validation of microfluidic-assisted biofabricated 3D constructs. a) Chick chorioallantoic membrane (CAM) model has been implemented using fertilized chick eggs and monitoring over time the development of vasculature surrounding (i) empty, (ii) factor-less controls, (iii) VEGF-gradient and (iv) VEGF-loaded 3D constructs. b) Vessel quantification produced (i) Chalkley score, (ii) area and (iii) diameter measurements. c) Subcutaneous implantation of 3D constructs was carried out over 4 weeks. d) Histological staining with hematoxylin & eosin (H&E) and Sirius red were carried out on (i, v) control, (ii, vi) TGFβ, (iii, vii) HBMSCs and (iv, viii) combination of HBMSCs and TGFβ. Statistical significance was calculated via one-way ANOVA. Results are expressed as mean ± SD of at least three replicates for each experiment,**p<0.01, *p<0.05. Scale bars: a, 10 mm; d,i-iv, 100 µm; d,v-vii, 50 µm;

To further explore the pre-clinical potential of the 3D constructs produced, functionally-graded 3D constructs with TGF-β1 and HBMSCs were engineered and implanted subcutaneously in mice to evaluate the potential for skeletal regeneration (**Figure 9c**). The graded patterning was here employed to fabricate a templating model for skeletal formation harnessing the synergistic effect of TGF-β1 and HBMSCs [61]. The histological evaluation with H&E (**Figure 9d-i-iv**) and Sirius red (**Figure 9d-v-viii**) staining on samples harvested after 4 weeks of implantation revealed that the inclusion of cells combined with graded TGF-β1 was found to elicit a superior collagenic matrix deposition compared to acellular and GF-free controls. This was also found consistent with previous findings [40,62], which demonstrated the ability of 3D bioprinted nanoclay-based materials to support skeletal regeneration *in ovo* and *in vivo*.

## Conclusions

The integration of microfluidic technologies with 3D bioprinting represents a major advancement in biofabrication, addressing critical limitations of conventional 3D bioprinting, such as poor cellular compartmentalization, material interface instability, and limited gradient biomaterial distribution. Current microfluidic-assisted technologies often fail to fabricate physiologically-relevant tissue interfaces and substitutes. In this study, we developed a novel microfluidic printhead to enhance 3D bioprinting by producing hierarchical 3D models of human tissues. This innovative approach not only streamlines microfluidic fabrication reducing manufacturing variability, but also could help to engineer increasingly hierarchical tissue interfaces for modelling and regeneration purposes. Here, we demonstrated the unparalleled ability to pattern different cell populations within a single 3D structure, thereby establishing distinct yet interconnected biological niches. A unique flow-focusing microfluidic printhead has been here employed to dynamically modulate fibre diameter and deposition, enabling tailored assembly of bioprinted constructs. By integrating a micromixing technology, the system could deliver distinct densities of cells and biomaterials, allowing independent patterning of multiple cell types. This would facilitate the engineering of compartmentalized micro-environments within the same 3D structure, simulating skeletal niches. Building on this, functionally-graded skeletal constructs were biofabricated resulting functional depending on the cellular density post-printing. The patterning of VEGF elicited a compartmentalized vascular ingrowth response, while TGF-β enhanced the differentiation of HBMSCs *in vivo*. These findings highlight the ability of the microfluidic-based system to control spatiotemporal deposition of biomaterials and cells, improving biomimicry and functionality of 3D skeletal constructs. The resulting multi-functional scaffolds, with compartmentalized skeletal microenvironments, open new avenues in tissue engineering, regenerative medicine, and drug discovery, advancing towards the engineering of increasingly physiologically relevant tissue interface models and potentially obviating the need for traditional animal testing procedures.

## Acknowledgements

GC acknowledges funding from ON Foundation (no. 22-214) and MTF Biologics (OSTEOMIMIC). This research was partially funded by grants from ERC-2019-Synergy Grant (ASTRA, n. 855923); EIC-2022-PathfinderOpen (ivBM-4PAP, n. 101098989); Project “National Center for Gene Therapy and Drugs based on RNA Technology” (CN00000041) financed by NextGeneration EU PNRR MUR—M4C2—Action 1.4—Call “Potenziamento strutture di ricerca e creazione di “campioni nazionali di R&S” (CUP J33C22001130001). The authors wish to thank the microscopy facility at Center for Life Nano- and Neuro-Science, Fondazione Istituto Italiano di Tecnologia. Cartoons in figures were created with BioRender.

## Notes on contributors

**Federico Serpe**: Writing – original draft, Visualization, Validation, Methodology, Investigation, Formal analysis.

**Lucia Iafrate**: Writing – review & editing, Methodology, Investigation.

**Marco Bastioli**: Writing – review & editing, Methodology, Investigation.

**Martina Marcotulli**: Methodology, Investigation.

**Caterina Sanchini**; Writing – review & editing, Methodology, Investigation.

**Valeria De Turris**: Methodology, Investigation,

**Michele D’Orazio**: Writing – review & editing, Methodology, Investigation.

**Biagio Palmisano**: Methodology, Investigation

**Arianna Mencattini**: Writing – review & editing, Methodology

**Eugenio Martinelli**: Resources, Funding acquisition

**Mara Riminucci**: Resources, Funding acquisition

**Carlo Massimo Casciola**: Resources, Funding acquisition

**Giancarlo Ruocco**: Resources, Funding acquisition

**Chiara Scognamiglio**: Conceptualization, Writing – review & editing, Supervision, Methodology, Project administration.

**Gianluca Cidonio**: Conceptualization, Writing – review & editing, Supervision, Visualization, Methodology, Project administration, Resources, Funding acquisition.

## Data availability statement

Data will be available upon reasonable request.

### Ethical statement

This study was performed in accordance with the Declaration of Helsinki. Human primary cell lines included in this study were approved as part of this study protocol. This human study was approved by Institutional Review Board (Department of Molecular Medicine, Sapienza University of Rome, Italy). All adult participants provided written informed consent to participate in this study. Animal study was approved by Institutional Review Board (Department of Molecular Medicine, Sapienza University of Rome, Italy). Animals had ad libitum access to standard mouse chow and water.

## Supplementary Information

**Movie S1.**
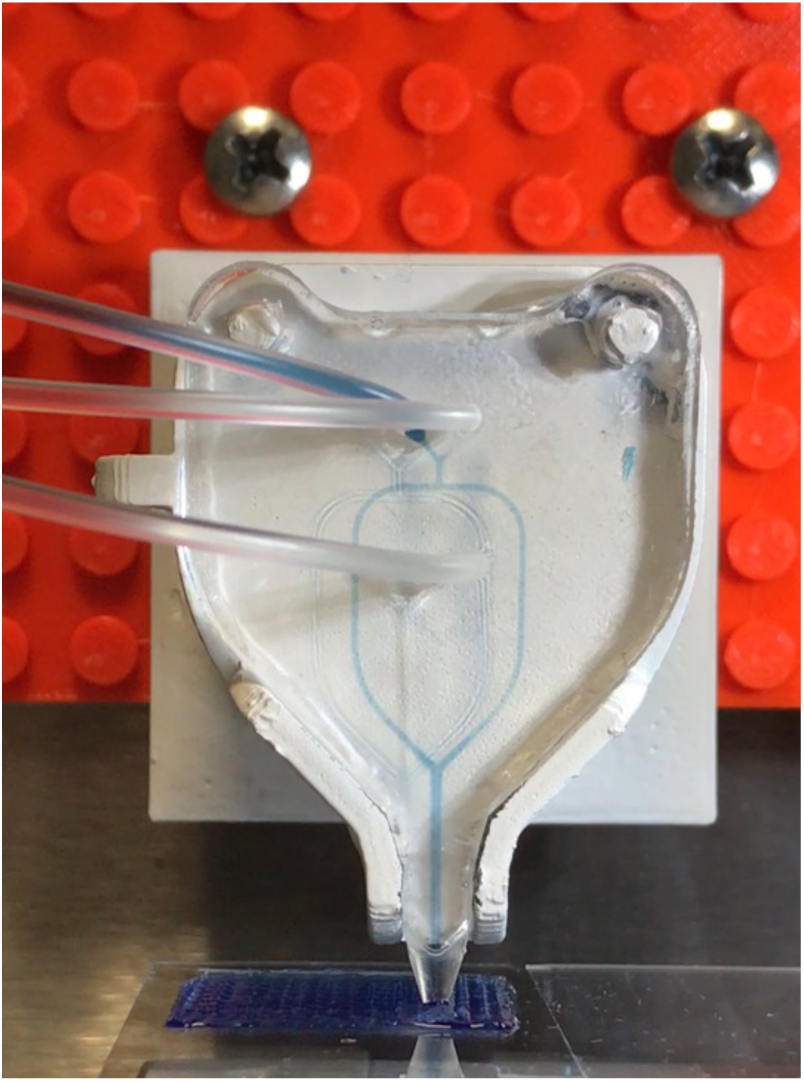
Video of active microfluidic-assisted 3D bioprinting during scaffolds fabrication

**Figure S1.**
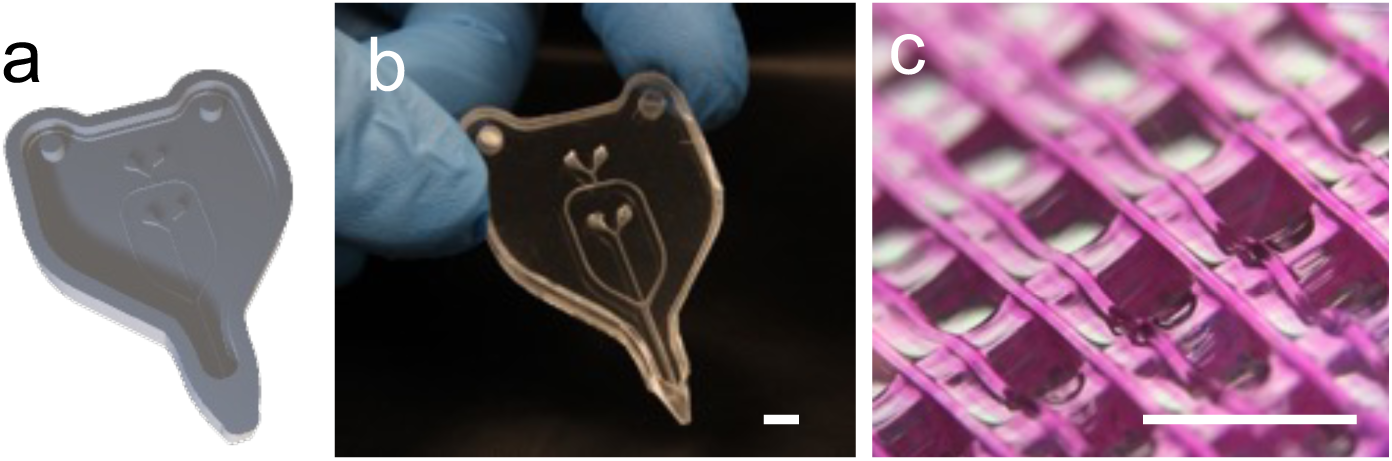
Microfluidic printhead. a) Render and b) macrograph of the engineered monolithic full-PDMS microfluidic printhead macrographs with cut-out supports for ease installation on 3D bioprinter, used to produce c) lattice scaffolds in 3D. Scale bars: 1 mm

**Figure S2.**
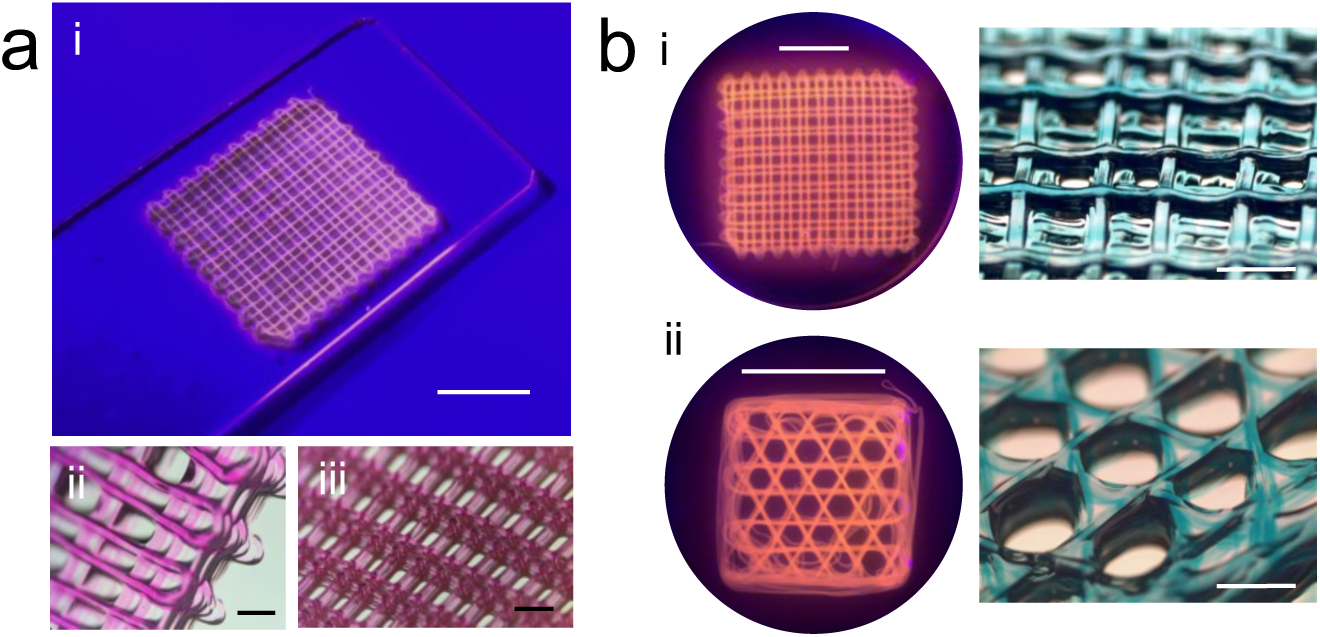
3D printed scaffold (a-i) with close-up on grid details showing the fabrication precision. b) Scaffolds with different infill geometries. On the left, top images of the entire 3D scaffolds, on the right, a magnified view of the infill pattern. Scale bars: a-i, 5 mm; a-ii,iii, 1 mm; b-i,ii (top view) 10 mm; b-i,ii (close-up view) 1 mm.

**Figure S3.**
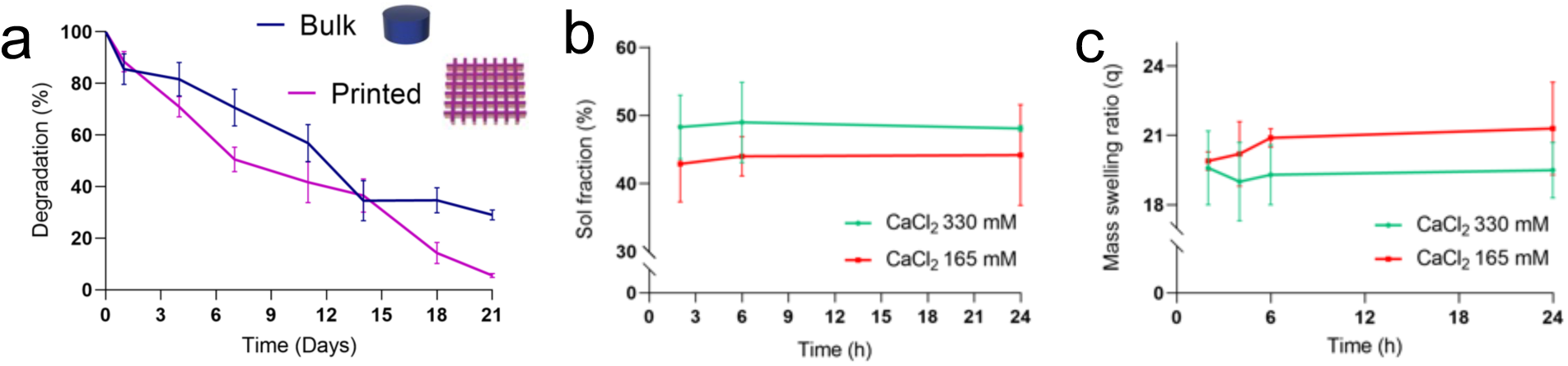
Characterization of 3D printed scaffolds. a) Comparison of the degradation rate between bulk and 3D printed scaffolds in 21 days. b) Comparison of the swelling properties of LAG crosslinked with different CaCl_2_ solutions. On the left, the percentage of solute fraction and on the right, the mass swelling ratio. Results are expressed as mean ± SD with n=3 replicates for each experiment.

**Figure S4.**
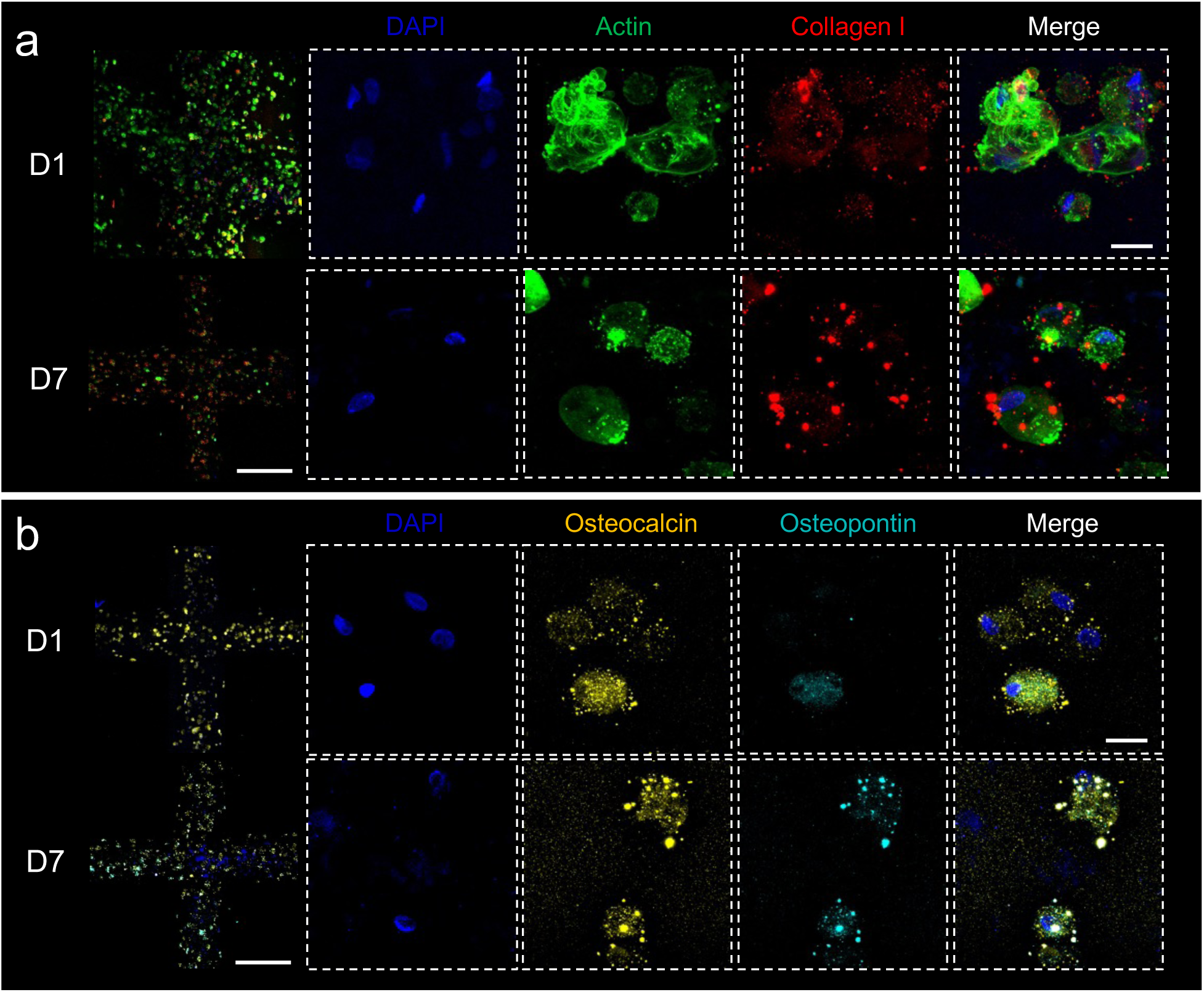
Immunofluorescence confocal imaging of microfluidic-assisted 3D bioprinting constructs stained to investigate a) morphological (DAPI, Actin, Collagen I) and b) skeletal (DAPI, Osteocalcin, Osteopontin) markers on HBMSCs. Scale bar: grid (500 μm), cells (20 μm)

