## Supplementary Info for "Engineering a microfluidic-assisted 3D bioprinting approach for the hierarchical control deposition and compartmentalisation of graded bioinks"

* Corresponding Authors:

### Supplementary Information

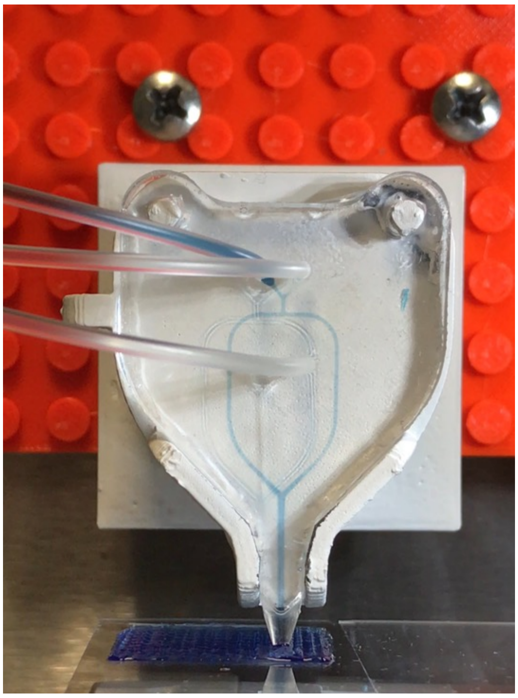

**Movie S1.** Video of active microfluidic-assisted 3D bioprinting during scaffolds fabrication

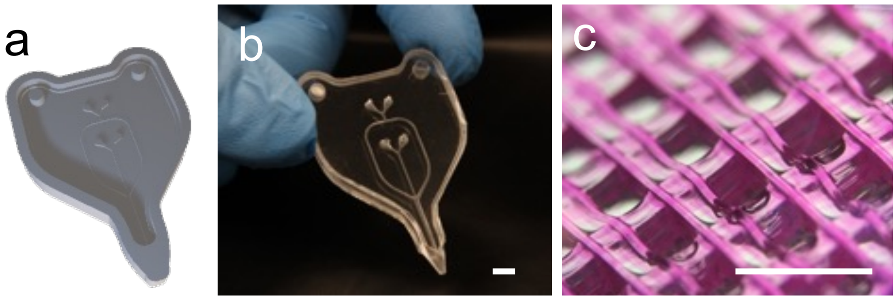


**Figure S1.** Microfluidic printhead. a) Render and b) macrograph of the engineered monolithic full-PDMS microfluidic printhead macrographs with cut-out supports for ease installation on 3D bioprinter, used to produce c) lattice scaffolds in 3D. Scale bars: 1 mm

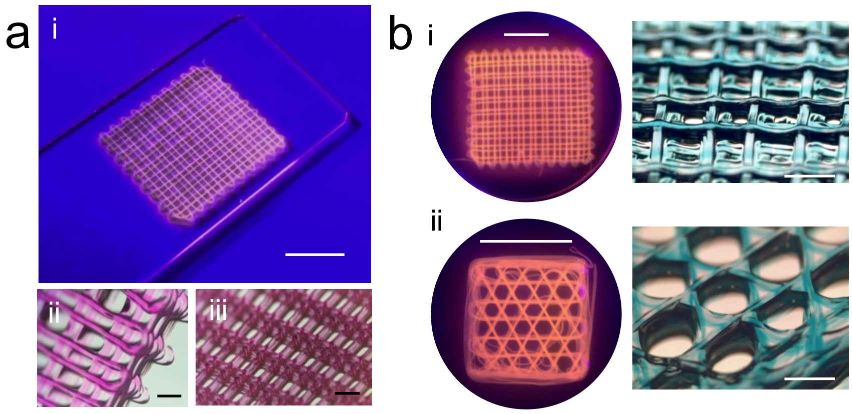


**Figure S2**. 3D printed scaffold (a-i) with close-up on grid details showing the fabrication precision. b) Scaffolds with different infill geometries. On the left, top images of the entire 3D scaffolds, on the right, a magnified view of the infill pattern. Scale bars: a-i, 5 mm; a-ii,iii, 1 mm; b-i,ii (top view) 10 mm; b-i,ii (close-up view) 1 mm.

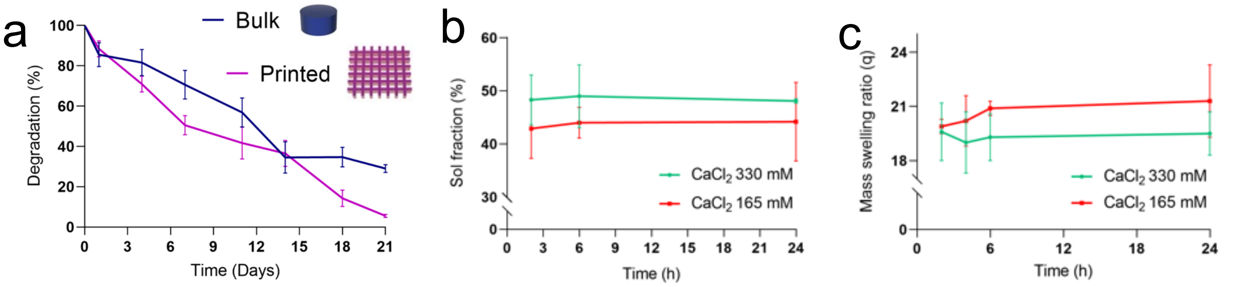


**Figure S3.** Characterization of 3D printed scaffolds. a) Comparison of the degradation rate between bulk and 3D printed scaffolds in 21 days. b) Comparison of the swelling properties of LAG crosslinked with different $CaCl_2$ solutions. On the left, the percentage of solute fraction and on the right, the mass swelling ratio. Results are expressed as mean ± SD with n=3 replicates for each experiment.

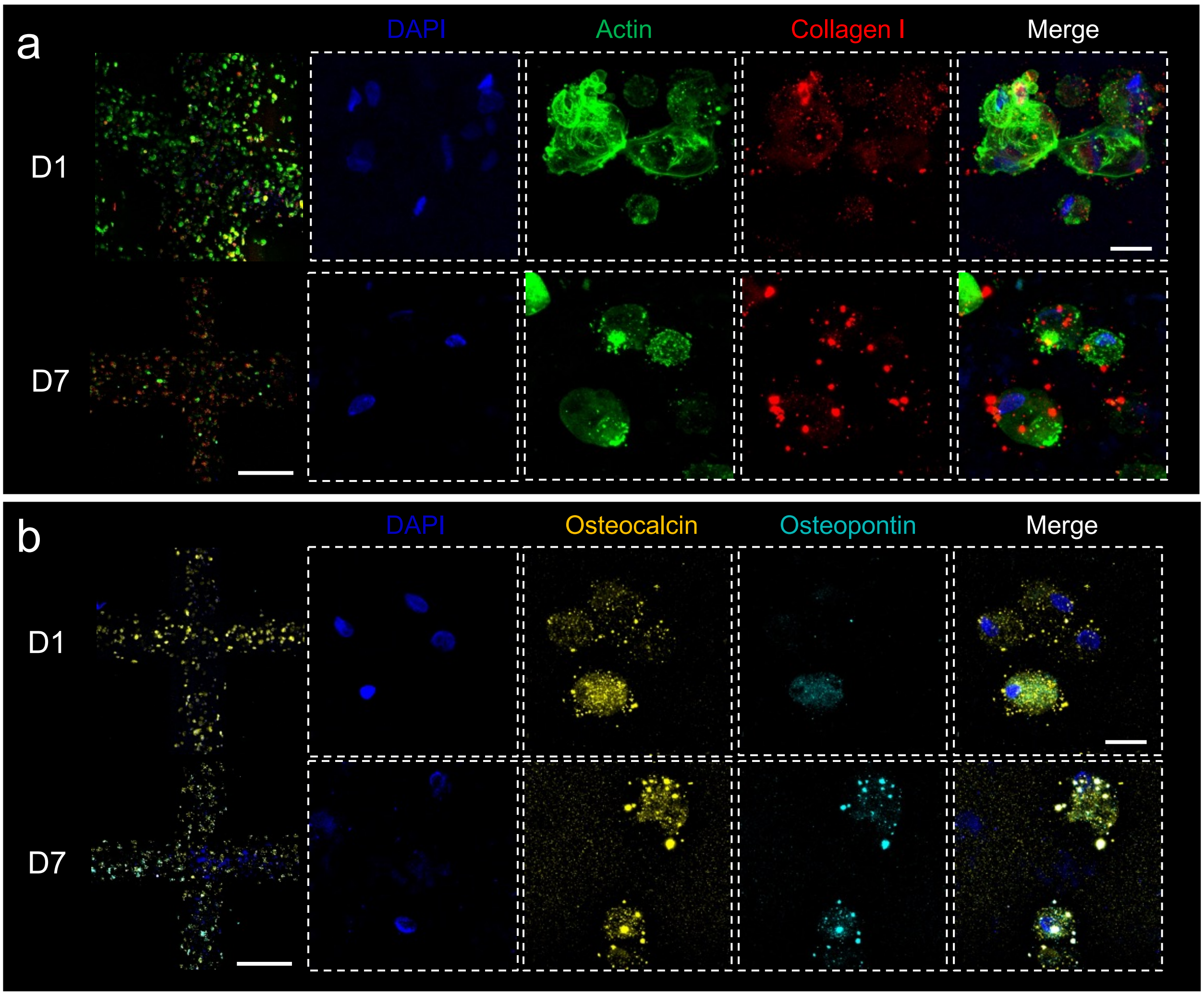


**Figure S4.** Immunofluorescence confocal imaging of microfluidic-assisted 3D bioprinting constructs stained to investigate a) morphological (DAPI, Actin, Collagen I) and b) skeletal (DAPI, Osteocalcin, Osteopontin) markers on HBMSCs. Scale bar: grid (500 μm), cells (20 μm)
